# Phosphate Starvation Signaling Increases Mitochondrial Membrane Potential through Respiration-independent Mechanisms

**DOI:** 10.1101/2022.10.25.513802

**Authors:** Yeyun Ouyang, Corey N. Cunningham, Jordan A. Berg, Ashish G. Toshniwal, Casey E. Hughes, Jonathan G. Van Vranken, Mi-Young Jeong, Ahmad A. Cluntun, Geanette Lam, Jacob M. Winter, Emel Akdoǧan, Katja K. Dove, Steven P. Gygi, Cory D Dunn, Dennis R Winge, Jared Rutter

## Abstract

Mitochondrial membrane potential directly powers many critical functions of mitochondria, including ATP production, mitochondrial protein import, and metabolite transport. Its loss is a cardinal feature of aging and mitochondrial diseases, and cells closely monitor membrane potential as an indicator of mitochondrial health. Given its central importance, it is logical that cells would modulate mitochondrial membrane potential in response to demand and environmental cues, but there has been little exploration of this question. We report that loss of the Sit4 protein phosphatase in yeast increases mitochondrial membrane potential, both through inducing the electron transport chain and the phosphate starvation response. Indeed, a similarly elevated mitochondrial membrane potential is also elicited simply by phosphate starvation or by abrogation of the Pho85-dependent phosphate sensing pathway. This enhanced membrane potential is primarily driven by an unexpected activity of the ADP/ATP carrier. We also demonstrate that this connection between phosphate limitation and enhancement of the mitochondrial membrane potential is also observed in primary and immortalized mammalian cells as well as in *Drosophila*. These data suggest that mitochondrial membrane potential is subject to environmental stimuli and intracellular signaling regulation and raise the possibility for therapeutic enhancement of mitochondrial functions even with defective mitochondria.

## Introduction

Mitochondria are a central hub for many cellular processes including ATP production, redox control, biosynthetic programs, and signaling (Chandel, 2015; Pagliarini & Rutter, 2013; Spinelli & Haigis, 2018). Each function—critical for cell homeostasis—rely on the ability of mitochondria to maintain a membrane potential across the inner membrane of this double-membrane organelle. This mitochondrial membrane potential (MMP, ΔΨm) directly provides the energy to power ATP synthesis, mitochondrial protein import, and metabolite and ion transport. Either directly or indirectly, it also provides signaling mechanisms to assist in adapting cellular behavior that can be critical to cell health.

Therefore, it is not surprising that impaired MMP is highly correlated with cellular dysfunction in aging (Hagen et al., 1997; C. E. Hughes et al., 2020; Leprat et al., 1990; Mansell et al., 2021; Sastre et al., 1996; Sugrue & Tatton, 2001) and a variety of diseases, including primary mitochondrial disease (Burelle et al., 2015; James et al., 1996) and heart failure (Cluntun et al., 2021; Sharov et al., 2005). While it is likely that MMP reduction plays a causal role in the pathogenesis of these diseases, the tools to formally test its impact on each disease are limited. In the context of aging, activating an artificial proton pump in *C. elegans* restores the loss of MMP typical of aging and is sufficient to extend lifespan (Berry et al., 2022), which suggests causality in this case. In addition, using the same manipulation to ectopically increase MMP improves the survival of *C. elegans* treated with electron transport chain inhibitors (Berry et al., 2020). These data raise the possibility that low MMP might cause pathology in the context of aging, and perhaps other diseases, and that strategies to restore membrane potential in cells might therefore be therapeutically transformative.

The canonical mechanism to generate MMP is by complexes I, III and IV of the electron transport chain (ETC), which pump protons from the mitochondrial matrix to the intermembrane space (IMS). This intricate process extracts high-energy electrons and passes them through the ETC complexes while using the resultant energy to pump protons from the matrix to IMS. The energy of these protons passing back to the matrix is then used to power ATP synthase, metabolite carriers, and protein translocases. However, several studies have described an alternative mechanism for the generation of MMP, namely ATP synthase running in reverse—hydrolyzing ATP to ADP and using the energy to pump protons to the IMS and augment the MMP (Junge & Nelson, 2015; Okuno et al., 2011). For example, Vasan, *et al*. reported that the MMP is maintained in complex III-deficient cells through this ATP synthase mechanism (Vasan et al., 2022). Such observations illustrate that cells will sacrifice hard-earned ATP to sustain their membrane potential, underlining the essentiality of MMP for cell well-being and insinuating the existence of control mechanisms to maintain MMP (Ernst et al., 2019; S. Liu et al., 2021; Martínez-Reyes et al., 2016).

MMP is required for viability and proliferation in most eukaryotic cells, but the strength of the MMP is highly variable between cells of different tissue origins and is dynamic across biological conditions (Huang et al., 2004; Mitra et al., 2009). For example, relative to normal cells, cancer cells tend to have a higher MMP (Davis et al., 1985; Heerdt et al., 2005; Summerhayes et al., 1982) as do cell experiencing amino acid starvation (Johnson et al., 2014). Generally, nutrient and other biological stress scenarios also modulate MMP (Hübscher et al., 2016, p. 70; Pan et al., 2011); in particular, oxidative stress has been shown to decrease MMP (Korshunov et al., 1997; Satoh et al., 1997). This heterogeneity in MMP amongst cell types and contexts led us to hypothesize that each cell might have an MMP setpoint that is tuned to the energetic and biosynthetic demands of the cell, and is perhaps responsive to nutrients and stressors in the environment. While there are well-appreciated negative consequences when MMP is too low, inappropriately elevated MMP can lead to toxic metabolic byproducts, such as reactive oxygen species. We only have sparse knowledge of whether cells actually have an MMP setpoint. If they do, how is it determined? What are the stimuli that are monitored to determine the setpoint? What are the signaling molecules that communicate this information? How is the machinery of mitochondrial bioenergetics altered to enact the setpoint and maintain this optimal MMP? Answering these questions will provide a much clearer understanding of the connection between cell physiology, mitochondrial bioenergetics, and human disease.

We became interested in MMP and its regulation through our previous studies of the mitochondrial fatty acid synthesis (mtFAS) system. We and others showed that loss of this system results in the absence of the lipoic acid cofactor as well as loss of acylated acyl carrier protein (ACP), which is required for the assembly and activation of many mitochondrial complexes, including each ETC complex (Angerer et al., 2017; Brody et al., 1997; Nowinski et al., 2020; Van Vranken et al., 2018). Using a genetic screen in yeast to identify genes required for the transcriptional alterations induced in mtFAS mutants, we found that the deletion of *SIT4* induces high MMP even in the absence of the ETC and ATP synthase. Building on this information, we identified genetic and environmental manipulations that increase MMP via ETC-dependent and independent mechanisms, including a non-canonical role for the ADP/ATP carrier. These results support the hypothesis that cells leverage available machineries to establish an MMP setpoint that is responsive to internal and external cues. We also identify signaling pathways and molecules that regulate this MMP setpoint. This study describes machinery involved in the modulation of MMP and provides effective tools to better understand the interplay between MMP and cellular health.

## Results

### *SIT4* deletion hyperpolarizes mitochondria

To understand the transcriptional reprogramming that occurs during the loss of mtFAS, and by extension, the interplay between dysfunctional mitochondrial and cell health, we re-analyzed an RNA-sequencing dataset (Berg et al., 2020) generated from a yeast mutant lacking mtFAS function (*mct1Δ*) (Schneider et al., 1997) before and after transitioning from glucose-to raffinose-containing medium––a manipulation that induces mitochondrial biogenesis. The canonical mitochondrial biogenesis transcriptional response was almost completely absent in the *mct1Δ* cells, as evidenced by the lack of induction of mRNAs encoding subunits of the ETC and ATP synthase (Fig. S1A). Instead, the *mct1Δ* mutant increased mRNA abundance of genes encoding proteins primarily related to mechanisms for acetyl-CoA production––a gene response signature that was not exhibited in wild-type cells (Fig. S1B). These data clearly indicate robust signaling from dysfunctional mitochondria to the nucleus, either to compensate or minimize the damage (Epstein et al., 2001; Garipler et al., 2014; Veatch et al., 2009). To better understand these mitochondria-to-nucleus transcriptional responses, we designed a genetic screen to identify the genes required for the transcriptional aberrations observed in *mct1Δ* cells. We selected one of the most upregulated genes (*CIT2*) to act as a reporter, and a gene with unchanged expression (*BTT1*) to act as a control. We integrated Firefly luciferase at the *CIT2* locus, Renilla luciferase at the *BTT1* locus, and the full-length *CIT2* and *BTT1* genes at the *HO* locus in the *mct1Δ* background (Fig. S1C). We screened through the non-essential gene deletion collection (~5,000 genes) (Giaever et al., 2002; Winzeler et al., 1999) and found 73 mutants (displayed in Fig. S1D) with a reduced ratio of expression from the native *CIT2* and *BTT1* promoters (i.e., reduced Firefly luciferase:Renilla luciferase ratio), suggesting an impaired *mct1Δ* mitochondrial dysfunction transcriptional signature.

We validated all 73 mutants by performing qRT-PCR on four additional genes that were upregulated in *mct1Δ* cells, *DLD3*, *ADH2*, *CAT2,* and *YAT1* (Fig. S1D). Some mutants still induced the expression of these genes, and others showed an absence of induction in only a subset of the four target genes (Fig. S1D). Of the mutants that reduced the abundance of all four transcripts, we focused on *SIT4* for two reasons. First, a *sit4Δ* mutant generated in our laboratory, as verification of this screening result, also displayed impaired induction of *CIT2*, *DLD3*, *ADH2*, and *CAT2* in response to *MCT1* deletion (Fig. 1A). Second, deletion of *SIT4* not only attenuated induction of these four genes, but also induced the expression of several genes encoding ETC subunits, including *QCR2* and *RIP1*, that were repressed in the *mct1Δ* mutant (Fig. 1B). Notably, deletion of *SIT4* alone––without a concomitant loss of *MCT1*––alters the abundance of these mRNAs tested, indicating that this transcriptional effect of *SIT4* deletion is independent of the underlying mitochondrial defects. These data suggest that *SIT4* is required for both the positive and negative transcriptional regulation elicited by mitochondrial dysfunction, and therefore we determined to define its role in this context.

**Figure 1:**
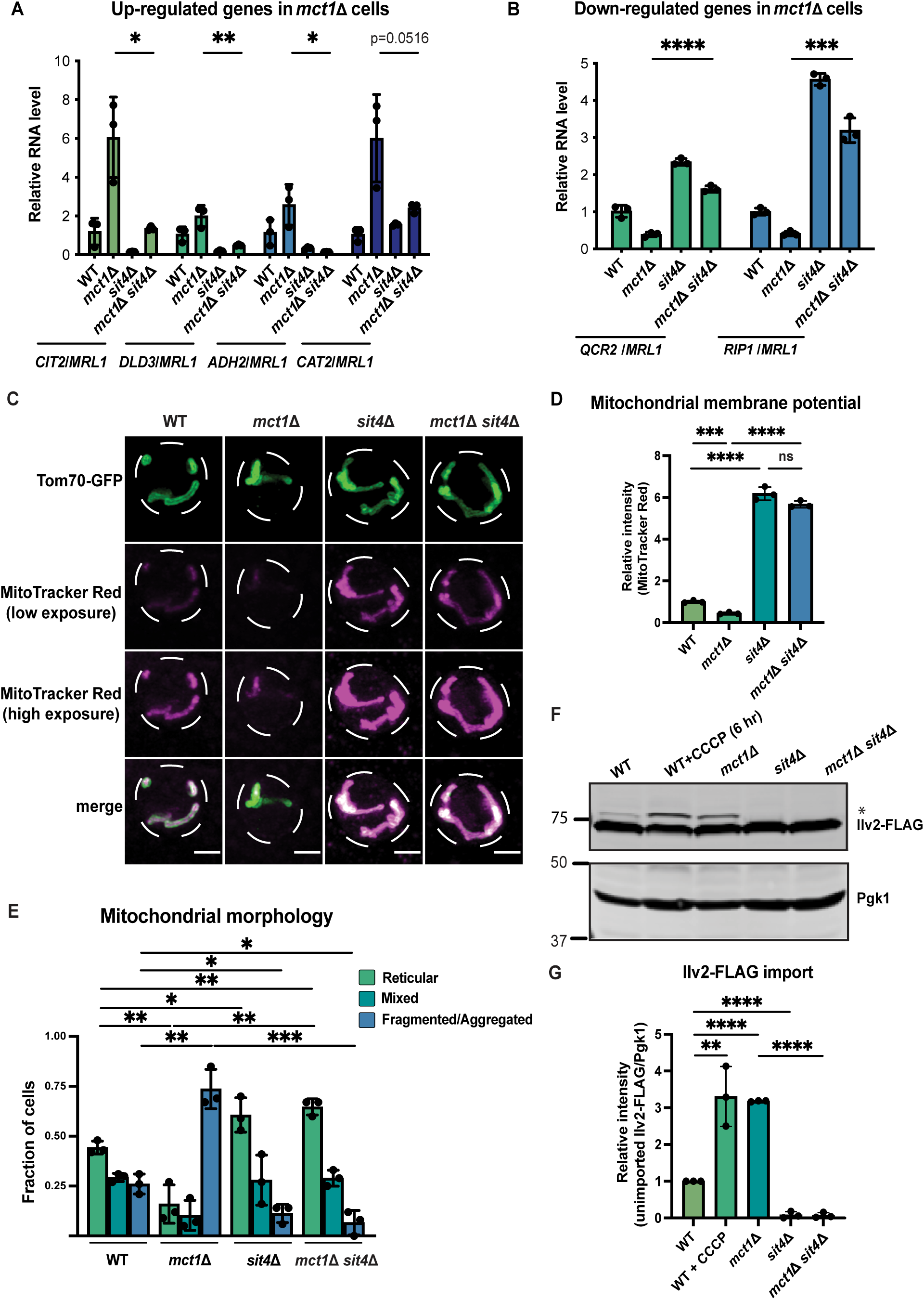
*sit4Δ* increases mitochondrial membrane potential in both wild-type and *mct1Δ* cells. (A and B) Normalized gene expression of *CIT2*, *DLD3*, *ADH2*, *CAT2*, *QCR2,* and *RIP1* measured three hours after switching from media containing 2% glucose as the sole carbon source to media containing 2% raffinose as the sole carbon source. Values were normalized to *MRL1*, a gene that was unchanged by deleting *MCT1* in our RNA-seq dataset. n = 3. Error bars represent the SD. Statistical significance was determined using an unpaired two-tailed t-test. *p ≤ 0.05; **p ≤ 0.005; ***p ≤ 0.0005; ****p ≤ 0.0001. (C) Representative images of wild-type (WT), *mct1Δ*, *sit4Δ*, and *mct1Δ sit4Δ* strains expressing Tom70-GFP from its endogenous locus stained with MitoTracker Red. Scale bar represents 2 μm. (D) Normalized mitochondrial membrane potential of wild-type (WT), *mct1Δ*, *sit4Δ*, and *mct1Δ sit4Δ* strains quantified by flow cytometry measurement of 10,000 cells stained with MitoTracker Red. n = 3. Error bars represent the SD. Statistical significance was determined using an unpaired two-tailed t-test. ns = not significant p > 0.05; ***p ≤ 0.0005; ****p ≤ 0.0001. (E) Quantification of the fraction of cells from wild-type (WT), *mct1Δ*, *sit4Δ*, and *mct1Δ sit4Δ* strains showing reticular, mixed, or fragmented/aggregated mitochondrial morphology. n = 3. Error bars represent the SD. Statistical significance was determined using an unpaired two-tailed t-test. *p ≤ 0.05; **p ≤ 0.005; ***p ≤ 0.0005. (F) Immunoblots of whole cell lysates extracted from wild-type (WT), *mct1Δ*, *sit4Δ*, and *mct1Δ sit4Δ* strains expressing Ilv2 endogenously tagged with FLAG. As a control, wild-type (WT) cells were treated with 25 μM CCCP for six hours. * indicates unimported Ilv2-FLAG. Pgk1 was immunoblotted as a loading control. Original immunoblots are displayed in Figure 1—source data. (G) Normalized quantification of (F). Import efficiency is the ratio of unimported Ilv2-FLAG (*) to Pgk1. n = 3. Error bars represent the SD. Statistical significance was determined using an unpaired two-tailed t-test. **p ≤ 0.005; ****p ≤ 0.0001. All original immunoblots used for quantification are displayed in Figure 1—source data.

*SIT4* is a serine/threonine phosphatase related to human PP6 that plays important roles in cell cycle regulation (Clotet et al., 1999; Fernandez-Sarabia et al., 1992), TOR signaling (Rohde et al., 2004; Torres et al., 2002), and tRNA modification (Abdel-Fattah et al., 2015). Additionally, a previous study by Garipler, *et al*. showed that deletion of *SIT4* in *rho^-^* cells, where the majority of mitochondrial DNA (mtDNA) is depleted, reverses some of the defects associated with mtDNA damage (Garipler et al., 2014). Despite their shared mitochondrial dysfunction, deletion of *MCT1* does not become *rho^-^* as shown by hybrid complementation assays (Fig. S1E).

We sought to understand how *SIT4* affects mitochondrial function and signaling upon loss of mtFAS. First, we measured the MMP in *sit4Δ* cells with or without deletion of *MCT1* using the membrane potential-dependent fluorescent dye, MitoTracker Red, which, when used at the appropriate concentration, accumulates in mitochondria in an MMP-dependent manner, such that the staining positively correlates with MMP. We used both microscopic imaging (Fig. 1C, S1F) and flow cytometry (Fig. 1D) as complementary methods to visualize and quantify MMP. With microscopic imaging, we were able to restrict the quantification of MitoTracker Red signal to that which co-localizes with mitochondria as marked by Tom70-GFP (Fig. 1C, S1F). Unexpectedly, we found that cells lacking *SIT4* alone exhibited very high mitochondrial membrane potential (Fig. 1C-D, S1F). The very high membrane potential observed in *sit4Δ* mutants was also maintained in *mct1Δ sit4Δ* double mutants (Fig. 1C-D, S1F). This was surprising given that *mct1Δ* cells lack assembly of the ETC, the major producer of MMP. The *mct1Δ sit4Δ* double mutant also exhibited a modest increase in mitochondrial area (Fig. S1G) and change in mitochondrial morphology. Therefore, we analyzed the localization pattern of Tom70-GFP and categorized the mitochondrial morphology of each cell as reticular, aggregated/fragmented, or mixed. We found that *mct1Δ* cells exhibited a more fragmented mitochondrial morphology, consistent with previous observations in other respiratory-deficient strains (Fig. 1E), whereas deletion of *SIT4* resulted in more reticular and less fragmented or aggregated mitochondria, both in the presence or absence of the *mct1Δ* mutation (Fig. 1E).

The import of most mitochondrial proteins from the cytosol depends upon and thus serves as a proxy for MMP. We used a yeast strain expressing a C-terminally FLAG-tagged Ilv2 from its endogenous locus (Dasari & Kölling, 2011). Upon import into mitochondria, the N-terminal mitochondrial targeting sequence (MTS) of Ilv2 is cleaved thereby allowing us to distinguish between imported and unimported species and providing a quantitative measurement of Ilv2 import as assessed by immunoblotting of whole cell lysates. In wild-type cells, the majority of Ilv2 is present as a lower molecular weight form with the MTS cleaved (Fig. 1F); however, a portion of the Ilv2 protein is visible as a higher molecular weight form suggesting it has not been imported into mitochondria and cleaved. Depletion of the MMP either by treatment for six hours with the ionophore CCCP or by deletion of *MCT1* reduced Ilv2 import and led to accumulation of the uncleaved protein (Fig. 1F-G). In contrast, deletion of *SIT4* caused complete mitochondrial protein import of Ilv2, whether or not *MCT1* was also deleted (Fig. 1F-G). These data demonstrate that loss of *SIT4* results in a mitochondrial phenotype suggestive of an enhanced energetic state: higher membrane potential, hyper-tubulated morphology and more effective protein import.

### *SIT4* deletion increases electron transport chain complex abundance

Because the changes in mitochondrial function observed in the *sit4Δ* mutant occur even in the mtFAS mutant background, which on its own creates a profound defect in mitochondrial respiratory complex assembly and energetics, we next asked how the deletion of *SIT4* increases MMP. RNA sequencing revealed that the *sit4Δ* mutant exhibited elevated expression of most of the genes encoding subunits of the ETC and ATP synthase, which could potentially promote ETC complex formation and function (Fig. 2A, Supplementary File 2). To directly assess the abundance of each ETC complex and supercomplexes, we performed blue-native PAGE (BN-PAGE) analysis on mitochondria isolated from wild-type, *mct1Δ*, *sit4Δ*, and, *mct1Δ sit4Δ* strains. This enables assessment of the assembly status of respiratory complex II, III, and IV, as well as the ATP synthase. Complex I is not included in our analysis due to its yeast homolog lacking the ability of proton pumping. As previously reported, mitochondria from *mct1Δ* mutant showed a complete loss of all the ETC complexes (Van Vranken et al., 2018) (Fig. 2B). Conversely and consistent with the RNA sequencing data, every ETC complex and supercomplex assembly was enriched in the *sit4Δ* mutant (Fig. 2B). Strikingly, deletion of *SIT4* also completely reversed the absence of and further enriched the abundance of ETC and ATP synthase complexes in *mct1Δ* cells. One potential explanation for this could be the restoration of mtFAS function in the *sit4Δ* mutant; however, *mct1Δ sit4Δ* cells still lacked acylated ACP (Fig. S2A) and lipoic acid as measured by lipoylation of Lat1 and Kgd2 (Fig. S2B), indicating that mtFAS remains inactive upon deletion of *SIT4*. Therefore, we conclude that deletion of *SIT4* enhanced the abundance of ETC complexes, even in the absence of a functional mtFAS pathway.

**Figure 2:**
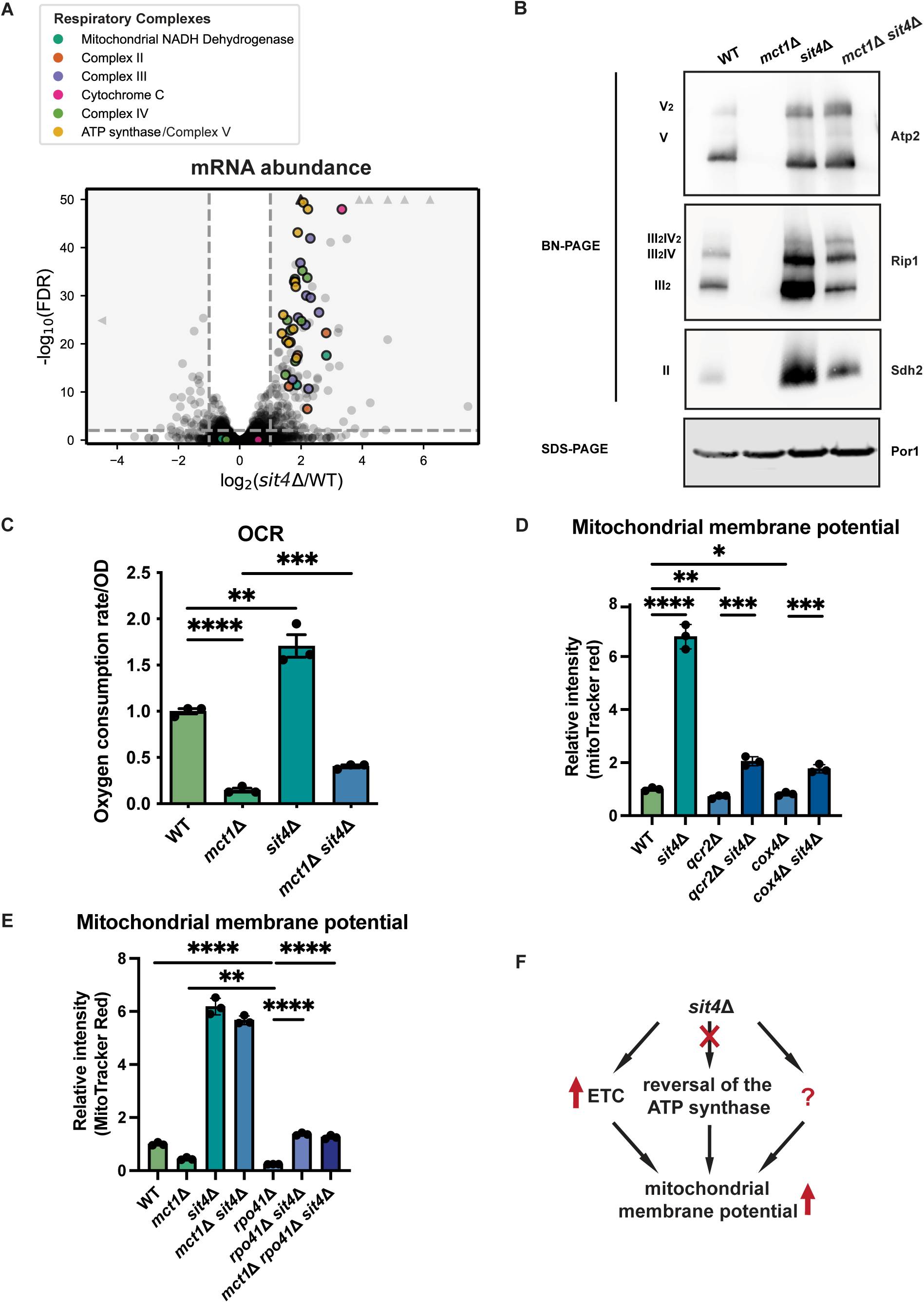
*sit4Δ* increases mitochondrial membrane potential through ETC-dependent and independent mechanisms. (A) Volcano plot of the transcriptomics data of *sit4Δ* vs. wild-type (WT). All genes encoding components of the ETC and ATP synthase that were detected by RNA sequencing are highlighted and color-coded. (B) Immunoblots of crude mitochondria extracted from wild-type (WT), *mct1Δ*, *sit4Δ*, and *mct1Δ sit4Δ* strains and separated on both BN-PAGE or SDS-PAGE. Membranes were blotted with indicated antibodies. Por1 was immunoblotted as a loading control. Original immunoblots are displayed in Figure 2—source data. (C) Normalized oxygen consumption rate (OCR) over optical density (OD) of the indicated strains grown in synthetic media containing 2% raffinose. n = 3. Error bars represent the SD. Statistical significance was determined using an unpaired two-tailed t-test. **p ≤ 0.005; ***p ≤ 0.0005; ****p ≤ 0.0001. (D and E) Normalized mitochondrial membrane potential of wild-type (WT), *sit4Δ*, *qcr2Δ*, *qcr2Δ sit4Δ*, *cox4Δ*, and *cox4Δ sit*4*Δ* strains quantified by flow cytometry measurement of 10,000 cells stained with MitoTracker Red. n = 3. Error bars represent the SD. Statistical significance was determined using an unpaired two-tailed t-test. ****p ≤ 0.0001. (F) Schematic of mechanisms through which *sit4Δ* increases mitochondrial membrane potential.

The enriched ETC complex abundance observed in the *sit4Δ* mutant is similar to what occurs duing glucose de-repression process (C. Jin et al., 2007). We therefore asked whether the increase of MMP and ETC complex assembly could be the result of impaired glucose repression. Normally, yeast cells activate a glucose repression program when grown in media with abundant glucose as a way of optimizing nutrient utilization. During this mode of growth, cells repress mitochondrial biogenesis and rely on glycolysis to provide the ATP required for rapid proliferation. We measured the MMP of wild-type and *sit4Δ* cells grown in glucose medium or in raffinose medium, which causes a loss of glucose repression. As expected from the increased expression of ETC genes, wild-type cells grown in raffinose-containing media had a higher MMP than glucose-grown cells (Fig. S2D). The *sit4Δ* mutant increased MMP to a similar degree in either growth medium (Fig. S2D). This result indicates that the increased MMP caused by *SIT4* deletion cannot be explained by glucose de-repression.

The presence of assembled ETC complexes in *sit4Δ* mutants raised the possibility that they are functional and contributing to MMP via proton pumping. To address this question, we measured the oxygen consumption rate (OCR) in all four yeast strains. As expected, the *mct1Δ* mutant exhibited a profound loss of OCR (Fig. 2C). On the other hand, *sit4Δ* cells exhibited an elevated OCR consistent with their increased abundance of ETC complexes. The *mct1Δ sit4Δ* cells showed an OCR that was increased relative to the *mct1Δ* single mutant, but still significantly lower than the wild-type OCR (Fig. 2C). Thus, we concluded that these ETC complexes are at least partially functional but are not sufficient to rescue the oxygen consumption defects found in the *mct1Δ* mutant. Consistent with a previous report (Jablonka et al., 2006), *sit4Δ* cells failed to grow on media containing a non-fermentable carbon source such as glycerol that requires mitochondrial respiration (Fig. S2C). This defect was rescued by re-expression of *SIT4* on a plasmid, confirming that *sit4Δ* cells have functional mtDNA. As expected, the *mct1Δ sit4Δ* double mutant also failed to grow under respiratory conditions (Fig. S2D). In conclusion, deletion of *SIT4* promotes the assembly of partially functional ETC complexes, but this is insufficient to rescue the respiratory defects of the *mct1Δ* mutant. Moreover, the observed modest increase in oxygen consumption is insufficient to explain the profound increase in MMP observed in the *mct1Δ sit4Δ* double mutant.

As a result of these data, we asked whether the enhanced MMP in *sit4Δ* mutants was dependent upon ETC proton pumping. We deleted subunits that are necessary for complex III (*QCR2*) and IV (*COX4*) assembly and function in both wild-type and *sit4Δ* mutant strains. Whether or not complex III or IV was inactivated, deletion of *SIT4* was sufficient to increase MMP (Fig. 2D). As mentioned previously, in the absence of a functional ETC, the mitochondrial ATP synthase has been shown to reverse direction and use the energy of ATP hydrolysis to pump protons across the mitochondrial inner membrane and generate a membrane potential. We treated wild-type cells and the *sit4Δ* mutant with oligomycin, an inhibitor of the F_o_ portion of the ATP synthase, to examine the directionality of the ATP synthase activity. Inhibiting the ATP synthase led to an increase of MMP in both wild-type and *sit4Δ* cells (Fig. S2E), suggesting that ATP synthase operates in the direction of MMP consumption and ATP generation in *sit4Δ* cells.

To orthogonally test whether the *sit4Δ* mutant can use mechanisms independent of ETC and ATP synthase, we obtained *rho^0^*cells that completely lack mtDNA and therefore have no functional ETC or ATP synthase. In spite of repeated efforts and success generating *rho^0^*cells in wild-type or other mutant backgrounds, we were unable to generate *rho^0^*cells in a *sit4Δ* strain. Therefore, we genetically eliminated the mitochondrial RNA polymerase, *RPO41*, which is required to transcribe all mtDNA-encoded transcripts (Greenleaf et al., 1986; Wang & Shadel, 1999). As a result, the *rpo41Δ* mutant lacks the proton-pumping ETC complexes III and IV as well as the protein-transducing F_o_ component of the ATP synthase (Fig. S2F). As expected, the MMP of the *rpo41Δ* mutant is very low, even significantly lower than that of *mct1Δ* cells (Fig. 2E). However, deletion of *SIT4* in the *rpo41Δ* mutant background restored the MMP to a level similar to wild-type cells (Fig. 2E). Intriguingly, complex II, which does not contain any mtDNA-encoded subunits, accumulated more assembled complex upon *SIT4* deletion (Fig. S2F). These results demonstrated that although ETC activity is required for the majority of the enhanced membrane potential observed in *sit4Δ* cells, *sit4Δ* mutants clearly leverage additional ETC- and ATP synthase-independent mechanisms to increase mitochondrial membrane potential (Fig. 2F).

### The *sit4Δ* mutant exhibits a phosphate starvation response

To discover the mechanisms through which the *sit4Δ* mutants generate MMP independent of the ETC or ATP synthase, we performed phosphoproteomics, as Sit4 is a protein phosphatase. Among the most enriched phosphoproteins, Pho84, Vtc3, and Spl2, are all involved in the regulation of intracellular phosphate levels (Figure 3A, Supplementary File 3). Pho84 is a high-affinity phosphate transporter on the plasma membrane, Vtc3 is involved in polyphosphate synthesis, and Spl2 mediates the downregulation of the low-affinity phosphate transporter during phosphate depletion. In addition, we cross-referenced our phosphoproteomics and RNA sequencing data (Fig. 2A, Supplementary File 2) and confirmed that many of the most significant transcriptional increases in the *sit4Δ* mutant were transcriptional targets of the PHO regulon (Fig. 3B), which stimulates phosphate acquisition triggered by either phosphate depletion or by dysregulation of the phosphate signaling pathway (Oshima, 1997; Paolo et al., 1997).

**Figure 3:**
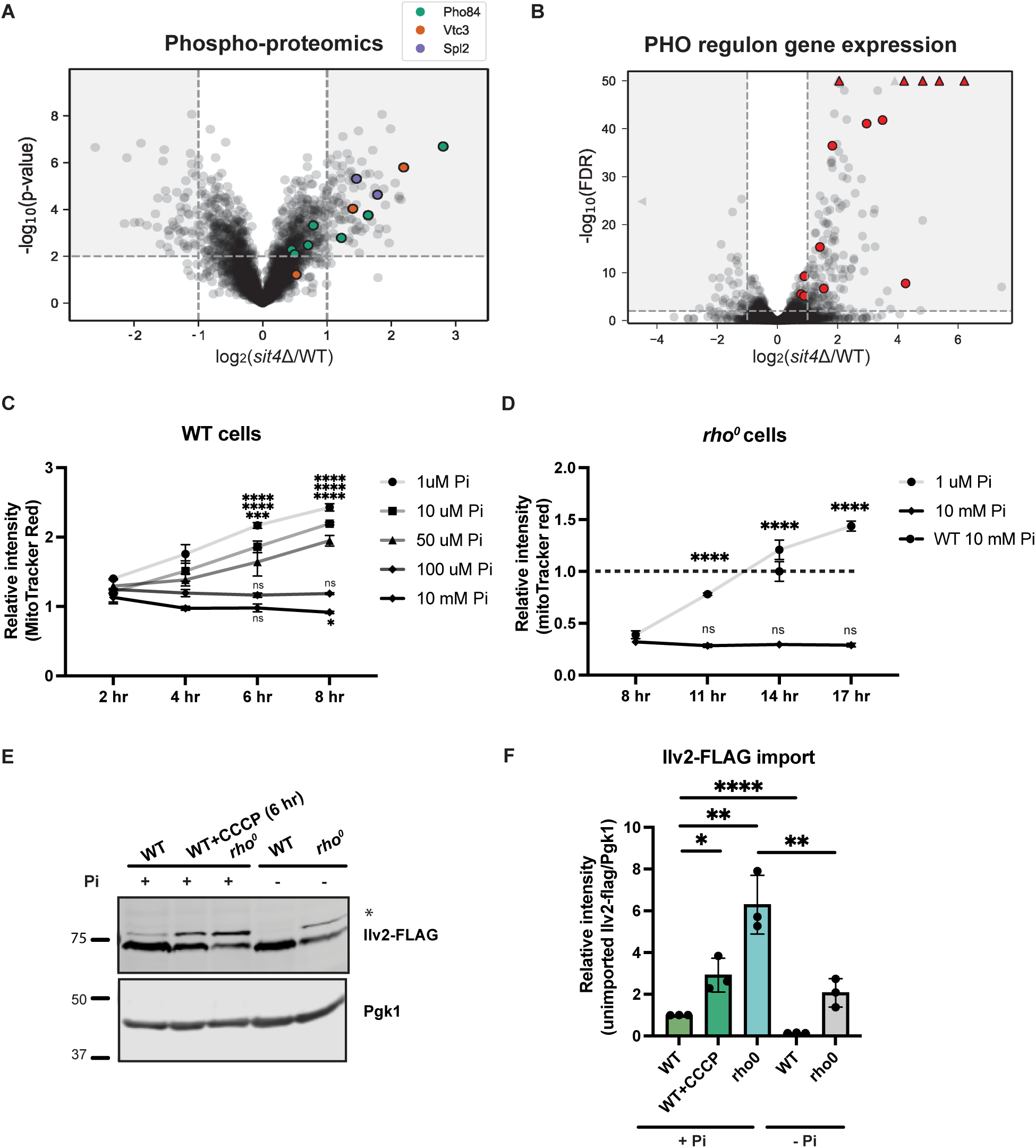
Phosphate starvation increases mitochondrial membrane potential through ETC-dependent and independent mechanisms. (A) Volcano plot of phospho-proteomics data of *sit4Δ* vs. wild-type (WT). Unique phosphorylation sites of Pho84 (green), Vtc3 (red), and Spl2 (purple) are highlighted. (B) Volcano plot of transcriptomics data of *sit4Δ* vs. wild-type (WT). All PHO regulon targets that are detected by RNA sequencing are highlighted in red. Triangle indicates that the −log10 (FDR) exceeds 50. (C and D) Normalized time course of mitochondrial membrane potential in wild-type (WT) and *rho^0^* strains measured by flow cytometry. The dashed line represents the membrane potential of wild-type (WT) cells grown in media containing 10 mM phosphate. n = 3. Error bars represent the SD. Statistical significance was determined using two-way ANOVA with Tukey’s multiple comparisons. ns = not significant p > 0.05; ***p ≤ 0.0005; ****p ≤ 0.0001. (E) Immunoblots of whole cell lysates extracted from wild-type (WT) or *rho^0^*cells expressing Ilv2 endogenously tagged with FLAG. Wild-type and *rho^0^*cells were grown in media containing either 10 mM of phosphate (+Pi) or 1 μM of phosphate (−Pi) for four hours or overnight, respectively. As a control, wild-type (WT) cells were treated with 25 μM CCCP for six hours. * indicates unimported Ilv2-FLAG. Pgk1 was immunoblotted as a loading control. Original immunoblots are displayed in Figure 3—source data. (F) Normalized quantification of (E). Import efficiency is the ratio of unimported Ilv2-FLAG (*) to Pgk1 band. n = 3. Error bars represent the SD. Statistical significance was determined using an unpaired two-tailed t-test. *p ≤ 0.05; **p ≤ 0.005; ****p ≤ 0.0001. All original immunoblots used for quantification are displayed in Figure 3—source data.

### Phosphate depletion increases mitochondrial membrane potential in wild-type and *rho^0^* <u>cells</u>

Given the unexpected activation of a phosphate starvation response upon deletion of *SIT4*, we tested whether modulation of environmental phosphate abundance could directly increase in MMP. All organisms acquire phosphate from the environment to build nucleic acids, phospholipids, and phosphorylated sugars and proteins. The synthetic media commonly used for growing yeast contains 7.5 mM inorganic phosphate. We depleted phosphate by growing yeast cells in media with close-to-normal phosphate levels (10 mM) or a series of lower phosphate concentrations (100, 50, 10, or 1 μM) and then measured the MMP using flow cytometry. Over an eight-hour time course, we observed an increase in the MMP of wild-type cells grown in low phosphate concentrations (50, 10, or 1 μM), but not in cells grown in either 100 μM or 10 mM phosphate (Fig. 3C). Having found that *sit4Δ* cells exhibited increased MMP even in the absence of the ETC and the F_o_ subunit of ATP synthase (Fig. 2E), we performed a similar phosphate depletion experiment in *rho^0^*cells, which lack all components of these two proton-pumping systems. Indeed, *rho^0^* cells exhibited elevated MMP in response to phosphate depletion (1 μM Pi), although the response was delayed compared to wild-type cells (Fig. 3D). We also confirmed this observation using fluorescence microscopy to capture the MMP and mitochondrial morphology after depleting phosphate in the media for four hours in wild-type cells and overnight in *rho^0^* cells (Fig. S3A). The quantification of the MitoTracker Red signal co-localized within Tom70-GFP agreed with the measurement of MMP using flow cytometry (Fig. S3B). At the same time, no significant change in mitochondrial area was observed by depleting phosphate from the media (Fig. S3C). Finally, the quantification of mitochondrial morphology suggested that phosphate deprivation does not induce massive changes in mitochondrial morphology in wild-type cells (Fig. S3D). However, the fragmented and aggregated mitochondria in *rho^0^* cells was partially rescued by phosphate depletion (Fig. S3D). Consistent with the MMP increase, mitochondrial protein import, as measured by Ilv2-FLAG cleavage, was enhanced by phosphate depletion in wild-type cells (Fig. 3E-F). *rho^0^*cells have a significant import deficiency, but this defect was mostly rescued by phosphate depletion as well (Fig. 3E-F).

Next, we examined respiratory complex assembly to understand further ETC and ATP synthase-related factors contributing to MMP. Using BN-PAGE, we found that in wild-type cells, phosphate depletion led to a modest enrichment in ETC complex abundance compared to cells grown in normal phosphate concentrations (Fig. 4A). However, depleting phosphate in *rho^0^* cells failed to rescue the absence of the complexes. Similar to the complex II enrichment in *rpo41Δ sit4Δ* cells (Fig. S2F), phosphate depletion also led to an accumulation in assembled complex II in *rho^0^* cells (Fig. 4A).

**Figure 4:**
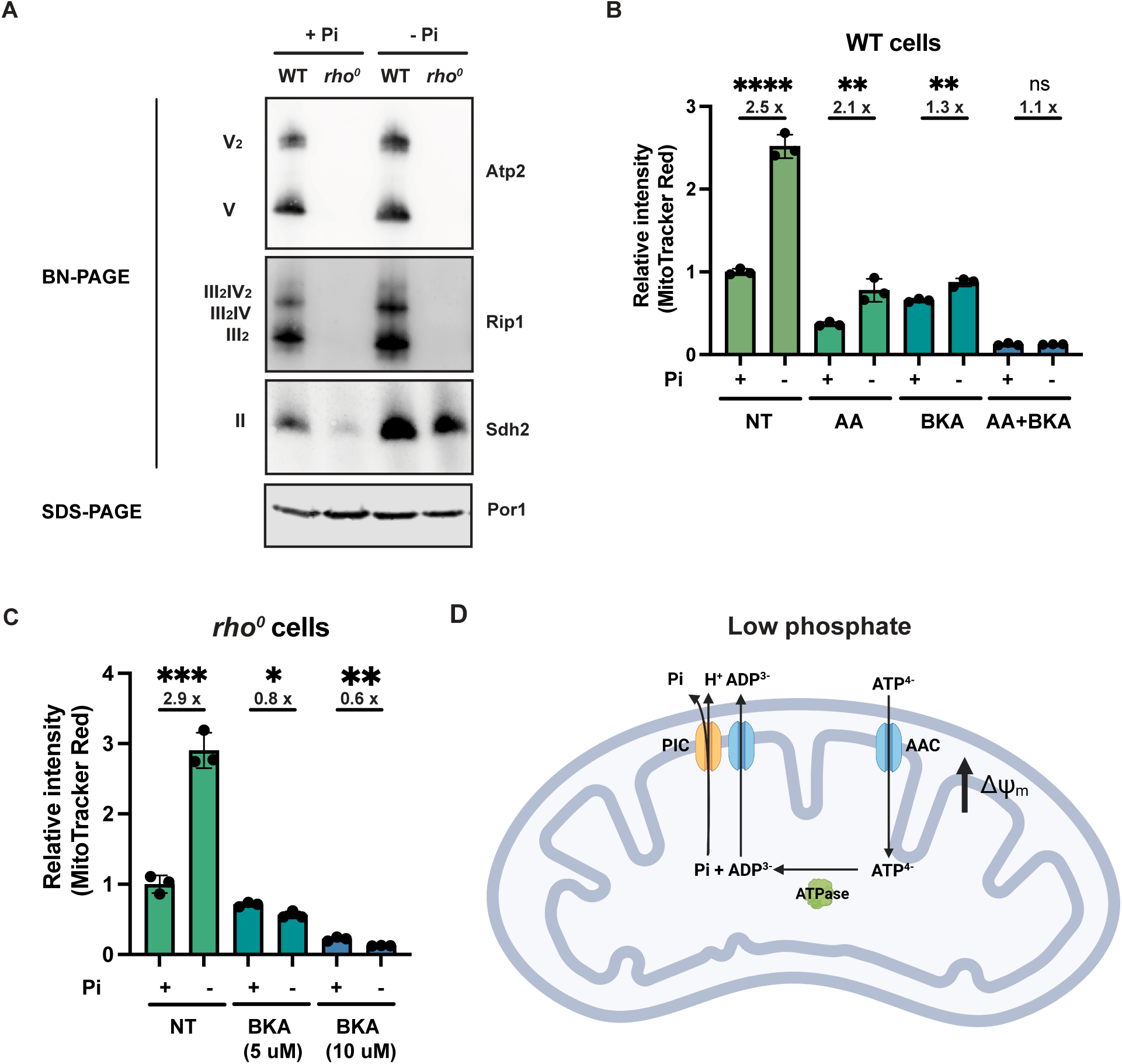
Phosphate depletion promotes mitochondrial membrane potential via ATP/ADP carrier in cells without ETC and ATP synthase. (A) Wild-type (WT) and *rho^0^* cells were grown in 10 mM (+Pi) or 1 μM (−Pi) phosphate-containing media for four hours or overnight, respectively. Crude mitochondria were extracted and separated by BN-PAGE or SDS-PAGE. Membranes were blotted with the indicated antibodies. Por1 was immunoblotted as a loading control. Original immunoblots are displayed in Figure 4—source data. (B) Wild-type (WT) cells were grown in 10 mM (+Pi) or 1 μM (−Pi) phosphate-containing media with or without drug treatment for four hours. Mitochondrial membrane potential was measured and quantified by flow cytometry. Error bars represent the SD. Statistical significance was determined using an unpaired two-tailed t-test. ns = not significant p > 0.05; **p ≤ 0.005; ****p ≤ 0.0001. (C) *rho^0^* cells were grown in 10 mM (+Pi) and 1 μM (−Pi) phosphate-containing media overnight and treated with or without bongkrekic acid (BKA) for four hours. Mitochondrial membrane potential was quantified by flow cytometry measurements of MitoTracker Red. Error bars represent the SD. Statistical significance was determined using an unpaired two-tailed t-test. *p ≤ 0.05; **p ≤ 0.005; ***p ≤ 0.0005. (D) Mechanisms of increased mitochondrial membrane potential induced by phosphate depletion.

We then treated cells with a series of mitochondrial inhibitors to parse out contributions of the ETC, ATP synthase, and other mechanisms to the MMP. Because electron transfer through complexes III and IV is tightly coupled to one another and with proton pumping, the complex III inhibitor antimycin A (AA) is sufficient to block the activities of both complexes. Most of the membrane potential was lost in cells treated with antimycin A in either normal or low phosphate-containing media (Fig. 4B); however, even in the presence of antimycin A, low phosphate still triggered a similarly fold elevation in membrane potential. Together with the BN-PAGE results, we conclude that ETC complexes are slightly enriched with phosphate depletion, but this is not required for the increase in MMP.

Bongkrekic acid is an inhibitor of the ADP/ATP carrier (AAC) (Lauquin & Vignais, 1976), which resides on the mitochondrial inner membrane and normally imports ADP and exports ATP to sustain mitochondrial ATP synthesis and cytosolic ATP consumption. Treatment of wild-type cells with bongkrekic acid significantly dampened the phosphate depletion-mediated increase in MMP (Fig. 4B). Importantly, combined treatment with both antimycin A and bongkrekic acid completely blocked the induction of MMP in response to low phosphate (Fig. 4B). As a genetic alternative to antimycin A inhibition of the ETC, we grew *rho^0^* cells, which have no complex III and IV nor complete ATP synthase, in low and high phosphate (Fig. 4C). As shown before, phosphate depletion triggers an enhanced MMP in *rho^0^*cells, but this is completely eliminated by bongkrekic acid in a dose-dependent manner (Fig. 4C).

These experiments suggest a mechanism whereby the depletion of phosphate increases MMP in an ETC- and ATP synthase-independent manner. When the ADP/ATP carrier imports ATP^4-^ and exports ADP^3-^, a net export of a positive charge occurs out of the matrix to the inter-membrane space (Fig. 4D). This activity must be coupled to ATP hydrolysis within the mitochondrial matrix by an as yet unidentified ATPase. It would also be coupled to the export of phosphate from the matrix, which is co-transported with a proton through the phosphate carrier. Our data suggest that when cells lack the proton pumping ability of the ETC––either by chemical (treatment with antimycin A) or genetic (loss of mtDNA in *rho^0^* cells) inhibition––and particularly during phosphate depletion, they instead rely on the ADP/ATP carrier to increase MMP to sustain critical mitochondrial functions.

### Phosphate starvation signaling induces mitochondrial membrane potential

The phosphate signaling system in yeast (Kaffman et al., 1994; Mouillon & Persson, 2006) employs the cyclin and cyclin-dependent kinase, Pho80 and Pho85, respectively. Under normal phosphate conditions, Pho85 is active but it is inhibited during phosphate depletion. When active, Pho85 phosphorylates and inactivates Pho4, a transcriptional factor that stimulates PHO regulon genes to promote phosphate acquisition, maintenance, and mobilization (Fig. S4A).

Deletion of *PHO85* in yeast cells results in constitutive activation of the PHO regulon even in a high phosphate environment. As a result, *pho85Δ* cells accumulate twice as much phosphate as wild-type cells (Gupta et al., 2019; N.-N. Liu et al., 2017). To elucidate how environmental phosphate starvation induces high MMP, we measured the MMP in *pho85Δ* mutants and found that, similar to *sit4Δ* cells or cells grown in phosphate-depleted media, MMP was significantly increased in *pho85Δ* cells (Fig. 5A, S4B-C). In the context of *MCT1* deletion, the *pho85Δ* also increased MMP, restoring the *mct1Δ* to near wild-type membrane potential (Figure A, S4B-C). Deletion of *PHO85* had no effect on mitochondrial area (Figure S4D). In spite of having an elevated MMP, the *pho85Δ* mutant shared a similar distribution of reticular, fragment/aggregated, or mixed mitochondrial morphology with wild-type cells (Fig. 5B). However, *PHO85* deletion normalized the fragmented and aggregated mitochondrial phenotype of *mct1Δ* cells (Fig. 5B). We also confirmed the MMP observation by demonstrating that the *pho85Δ* mutant had improved import of Ilv2-FLAG than wild-type cells, and *PHO85* deletion restored the import activity of the *mct1Δ* mutant to around wild-type levels (Fig. 5C-D). These phenotypes of the *pho85Δ* mutant strain suggest that the mechanism underlying the MMP increase in response to phosphate starvation is unlikely to be a direct result of intracellular phosphate insufficiency. Rather, these data support the hypothesis that the perception of phosphate starvation, and activation of the phosphate signaling response, is the key driver of MMP enhancement.

**Figure 5:**
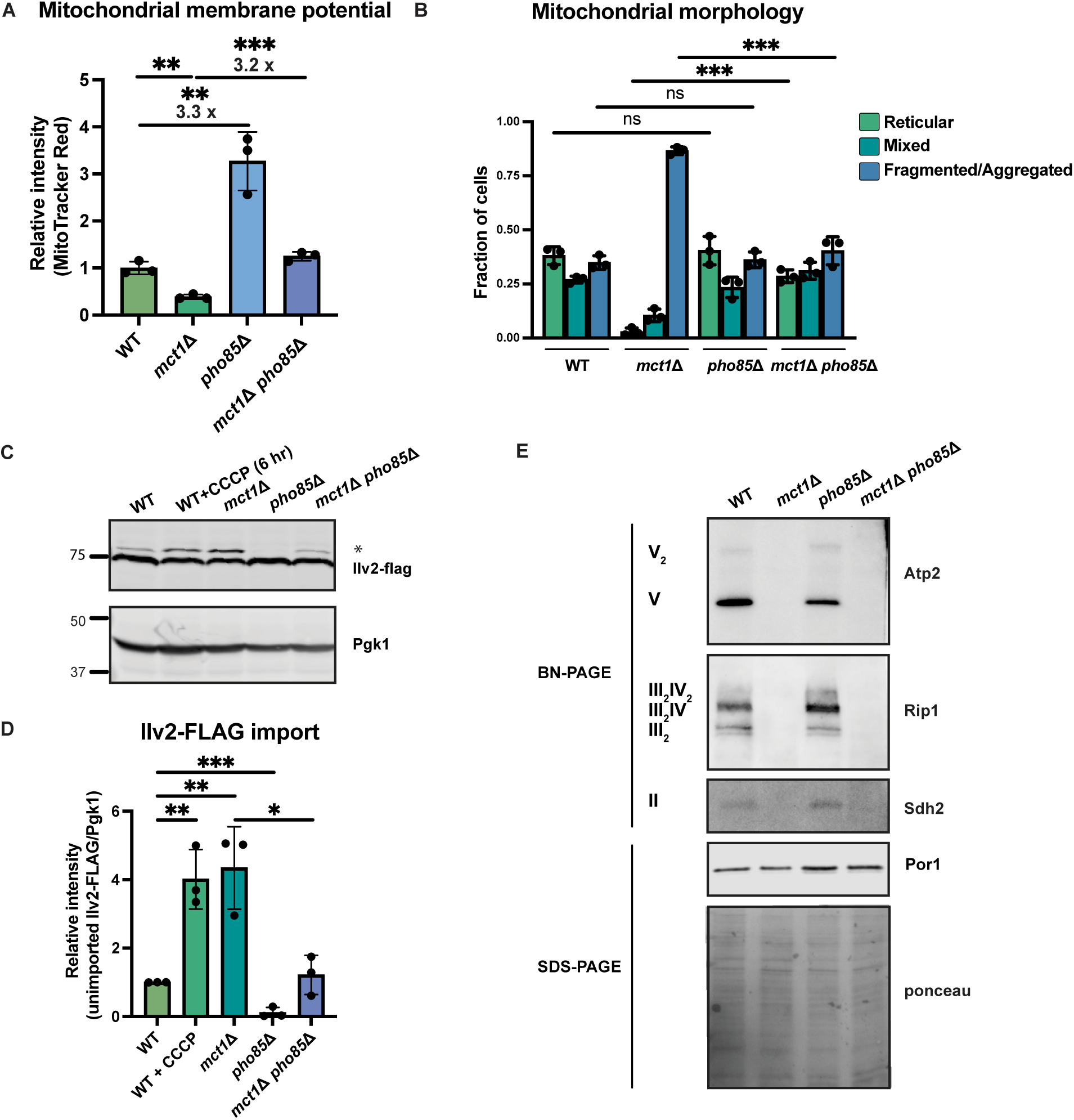
Activation of phosphate signaling increases mitochondrial membrane potential. (A) Normalized mitochondrial membrane potential of wild-type (WT), *mct1Δ*, *pho85Δ*, and *mct1Δ pho85Δ* strains quantified by flow cytometry measurement of 10,000 cells stained with MitoTracker Red. n = 3. Error bars represent the SD. Statistical significance was determined using an unpaired two-tailed t-test. **p ≤ 0.005; ***p ≤ 0.0005 (B) Quantification of the fraction of cells from wild-type (WT), *mct1Δ*, *pho85Δ*, and *mct1Δ pho85Δ* strains showing reticular, mixed, or fragmented/aggregated mitochondrial morphology. n = 3. Error bars represent the SD. Statistical significance was determined using an unpaired two-tailed t-test. ns = not significant p > 0.05; ***p ≤ 0.0005. (C) Immunoblots of whole cell lysates extracted from wild-type (WT), *mct1Δ*, *pho85Δ*, and *mct1Δ pho85Δ* cells expressing Ilv2 endogenously tagged with FLAG. As a control, wild-type (WT) cells were treated with 25 μM CCCP for six hours. * indicates unimported Ilv2-FLAG. Pgk1 was immunoblotted as a loading control. Original immunoblots are displayed in Figure 5—source data 1. (D) Normalized quantification of (C). Import efficiency is the ratio of unimported Ilv2-FLAG (*) to Pgk1. n = 3. Error bars represent the SD. Statistical significance was determined using an unpaired two-tailed t-test. *p ≤ 0.05; **p ≤ 0.005; ***p ≤ 0.0005. All original immunoblots used for quantification are displayed in Figure 5—source data 1. (E) Immunoblots of crude mitochondria extracted from wild-type (WT), *mct1Δ*, *pho85Δ*, and *mct1Δ pho85Δ* cells and separated by BN-PAGE or SDS-PAGE. Membranes were blotted with indicated antibodies. The membrane was stained with Ponceau S as a loading control. Original immunoblots are displayed in Figure 5—source data 2.

Finally, we asked whether deletion of *PHO85* could rescue the ETC complex assembly defect in the *mct1Δ* mutant, as observed in the *sit4Δ* mutant, which could be an explanation for the enhanced MMP. Consistent with phosphate depleted wild-type cells as shown in Figure 4A, there was a modest enrichment of ETC complexes in the *pho85Δ* mutant (Fig. 5E). In the context of the *mct1Δ* mutant strain, however, deletion of *PHO85* had no effect on the complete loss of ETC complexes. We reasoned that this lack of rescue was different from *SIT4* deletion because *pho85Δ* cells exhibit no increase in the abundance of mRNAs encoding ETC and ATP synthase subunits (Fig. S4E, compared to Fig. 2A for *sit4Δ*, Supplementary File 2). We therefore conclude that activating phosphate starvation signaling is sufficient to establish an elevated mitochondrial membrane potential, in a manner that is mostly independent of the ETC. In particular, *mct1Δ pho85Δ* cells exhibit a much higher MMP compared to the *mct1Δ* mutant, and this appears to largely be mediated by ETC- and ATP synthase-independent mechanisms.

### Depleting phosphate increases mitochondrial membrane potential in higher eukaryotes

Given the importance of mitochondrial membrane potential for human health and disease, we tested whether phosphate depletion might also enhance MMP in the HEK293T (embryonic kidney) and A375 (melanoma) human cells. Both HEK293T and A375 cells exhibited increased MMP after three days of growth in phosphate-free medium (Fig. 6A, S5A). We quantified the morphology of the mitochondrial network based on two parameters: summed branch length mean, which describes the mean of summed length of mitochondrial tubules in each independent structure, and network branch number mean which describes the mean number of attached mitochondrial tubules in each independent structure. Consistent with the increased MMP, HEK293T and A375 cells grown in low phosphate exhibited a more connected and elongated mitochondrial network (Fig. 6B, S5B). As an alternative strategy to limit phosphate uptake from the media, we treated these same cell lines with the phosphate transporter inhibitor phosphonoformic acid (PFA) for 48 hours. We observed a dose-dependent increase in MMP in both cell lines (Fig. 6C). We concluded that phosphate depletion induces a higher MMP in cultured mammalian cell lines. Immortalized cell lines may have adaptations that are not representative of native cells, so we decided to determine whether primary mammalian cells might also exhibit this phenomenon. Due to the special media requirements for culturing primary hepatocytes and the lack of commercially available phosphate-depleted media, we treated primary hepatocytes with PFA for 24 hours to generate a phosphate-depleted intracellular state. Similar to the immortalized cell lines, MMP increased in a dose-dependent manner (Fig. 6D).

**Figure 6:**
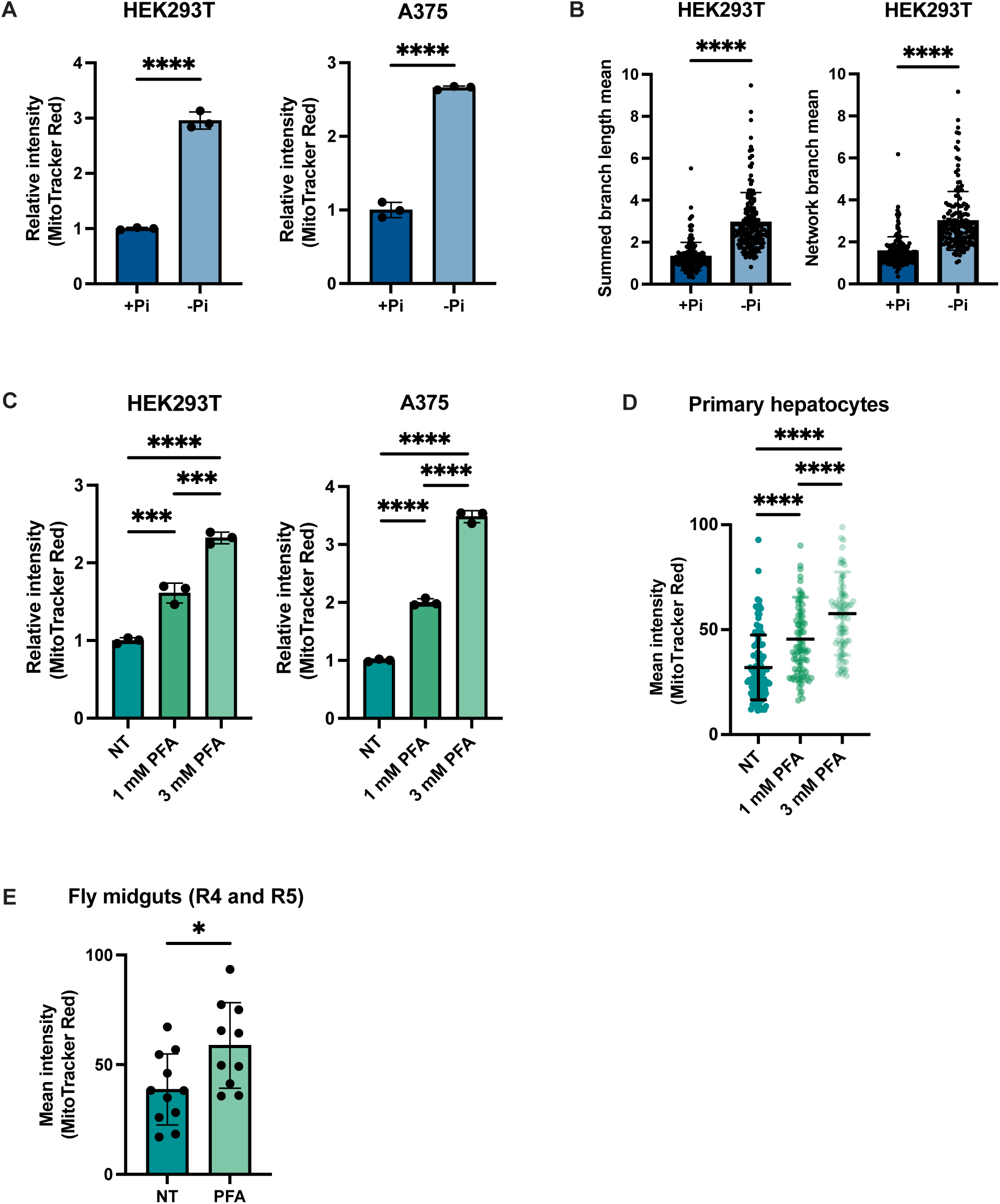
Phosphate depletion induces increased mitochondrial membrane potential in higher eukaryotic cells. (A and B) The indicated cell lines were cultured with 1 mM (+Pi) or no phosphate (−Pi) for three days. Mitochondrial membrane potential was quantified by flow cytometry measurement of 10,000 cells stained with MitoTracker Red. n = 3. Error bars represent the SD. Statistical significance was determined using an unpaired two-tailed t-test. ****p ≤ 0.0001. (C) Summed branch length mean and network branch mean were measured and calculated by Mitochondrial Network Analysis (MiNA). n = 3. Error bars represent the SD. Statistical significance was determined using an unpaired two-tailed t-test. ****p ≤ 0.0001 (D) HEK293T cells were treated with 0, 1, or 3 mM phosphonoformic acid (PFA) for 48 hours. Mitochondrial membrane potential was quantified by flow cytometry measurement of 10,000 cells stained with MitoTracker Red. n = 3. Error bars represent the SD. Statistical significance was determined using a one-way ANOVA with Tukey’s multiple comparisons. ***p ≤ 0.0005; ****p ≤ 0.0001. (E) Primary hepatocytes were treated with 0, 1, or 3 mM PFA for 24 hours, and then stained with MitoTracker Red and imaged. The mean intensity of mitochondrial membrane potential was quantified by measurement of by the MitoTracker Red fluorescent signal from ~80 cells per condition. Error bars represent the SD. Statistical significance was determined using a one-way ANOVA with Tukey’s multiple comparisons. ****p ≤ 0.0001. (F) 3-week-old flies from the control group or from the experiment group treated with 1 mM phosphonoformic acid (PFA) for 2 weeks were dissected. Their midguts (R4 and R5 region) were stained with TMRE and the fluorescent signal was quantified by microscopic imaging. Error bars represent the SD. Statistical significance was determined using an unpaired two-tailed t-test. *p ≤ 0.05.

Finally, we used the fruit fly *Drosophila melanogaster* to assess the relationship of phosphate starvation and MMP in an intact living animal. PFA has been previously used in the fly to generate a phosphate-depleted state (Bergwitz et al., 2013). PFA treatment is lethal during the larval stage in the fly developmental cycle, but can extend lifespan when administered to adult flies (Bergwitz et al., 2013). Indeed, PFA-treated adult flies exhibited a higher MMP than the control group as measured by quantification of TMRE-stained images of the fly midgut. We conclude, therefore, that the relationship between phosphate deprivation and mitochondrial membrane potential extends across evolution and is recapitulated in vitro and in vivo, suggesting its fundamental importance to eukaryotic biology.

## Discussion

The work described herein was initially intended to define the signaling and transcriptional network that underlies the positive and negative gene expression effects of a mutant that lacks the mtFAS system, and therefore lacks assembly of the respiratory system. This line of inquiry led to a series of observations that demonstrated a previously unappreciated role of environmental sensing and cellular signaling in controlling MMP. As a result, we propose the model that, based on energetic demand, environmental status, and intracellular signaling, cells establish and maintain a MMP “setpoint”, which is tailored to maintain optimal mitochondrial function. We find that cells deploy multiple ETC-dependent and -independent strategies to maintain that setpoint. Critically, we find that cells often prioritize this MMP setpoint over other bioenergetic priorities, even in challenging environments, suggesting an important evolutionary benefit.

We first made the observation that deletion of the *SIT4* gene, which encodes the yeast homologue of the mammalian PP6 protein phosphatase, normalized many of the defects caused by loss of mtFAS, including gene expression programs, ETC complex assembly, mitochondrial morphology, and especially MMP (Fig. 1). A previous study (Garipler et al., 2014) reported that the deletion of *SIT4* increased MMP in *rho*^-^ cells, although the mechanism underlying the phenomenon was not defined. We show herein that increased MMP in a *sit4Δ* mutant is independent of mtDNA damage and occurs in an otherwise wild-type strain (Fig. 1C-D). The mechanism whereby *SIT4* deletion elicits these mitochondrial effects seems to involve the induction of two gene expression programs: mitochondrial electron transport chain biogenesis and the phosphate starvation response. Through these effects, and perhaps other mechanisms, the *sit4Δ* mutant rescues the ETC assembly failure caused by loss of mtFAS (Fig. 2B). Indeed, the ETC complex abundance and MMP of an *mct1Δ sit4Δ* double mutant is substantially enhanced relative to a wild-type strain. This has important implications for our understanding of how mtFAS supports, but is not essential for, ETC assembly.

The transcriptional and phosphoproteomic effects of *SIT4* deletion (Fig. 3A-B) led us to the observation that phosphate deprivation, or the perception of phosphate starvation by elimination of the Pho85-dependent phosphate sensing system, was sufficient to increase MMP. Unlike phosphate starvation, the *pho85Δ* mutant has elevated intracellular phosphate concentrations. This suggests that the phosphate effect on MMP is likely to be elicited by cellular signaling downstream of phosphate sensing rather than some direct effect of environmental depletion of phosphate on mitochondrial energetics.

One of the more surprising findings from this work is that the phosphate starvation response increases MMP independently of either of the two well-established MMP generation mechanisms, proton pumping by the electron transport chain or ATP hydrolysis-dependent proton pumping by the ATP synthase. Phosphate starvation or the deletion of *PHO85* has minimal effects on the assembly of the respiratory complexes and does not restore detectable complexes to an *mct1Δ* strain, but still increases MMP. Phosphate deprivation even increases MMP in a mutant lacking the entire mitochondrial genome, and hence has no assembled ETC complex III or IV and no ATP synthase. Instead, our data suggest that the ADP/ATP carrier is utilized to increase MMP in response to phosphate starvation signaling (Fig. 4B-C). The import of ATP^4-^ produced by glycolysis, hydrolysis to ADP^3-^ and inorganic phosphate, and export of ADP^3-^ through the AAC, is an electrogenic process that is sufficient to sustain or enhance the MMP. Meanwhile, the inorganic phosphate released during ATP hydrolysis is exported with a proton through the phosphate carrier, which is an electroneutral process but contributes to the proton gradient. It was reported previously that *rho^0^* cells rely on the reverse transport of ATP and ADP through the ADP/ATP carrier in conjunction with ATP hydrolysis in the mitochondrial matrix to generate a minimal MMP (Appleby et al., 1999; Buchet & Godinot, 1998; X. J. Chen & Clark-Walker, 2000; Dupont et al., 1985; Kováčová et al., 1968). We now show that intracellular signaling can lead to an increased MMP even beyond the wild-type level in the absence of mitochondrial genome.

The elevated MMP setpoint has significant functional consequences, as demonstrated by the partially mitochondrial Ilv2 protein becoming completely imported and cleaved upon loss of Sit4 or phosphate depletion. It is likely that many other proteins either gain or enhance their mitochondrial matrix localization and activity as well. It has been previously described that changes in MMP can alter the import properties of proteins and thereby trigger cellular signaling events related to mitophagy, transcriptional responses, and others (Becker et al., 2012.; Berry et al., 2021.; S. M. Jin et al., 2010; Miceli et al., 2012; Rolland et al., 2019). The scope of responses elicited in cells experiencing high MMP, however, has not been previously interrogated.

What is the evolutionary advantage of hydrolyzing ATP to ADP and phosphate, while using the energy to increase the MMP? We propose a speculative hypothesis that this mechanism enables the liberation of needed phosphate from the most abundant labile store of phosphate, ATP. However, cleaving the phosphodiester bonds of ATP releases energy in addition to releasing phosphate. Using this mitochondrial mechanism enables the cell to capture that energy in the form of the MMP rather than having it be simply lost as heat. As a result, the elevated MMP is able to fuel mitochondrial processes and empowers the cell to better combat nutrient scarcity. This phenomenon also appears to be evolutionarily conserved as cellular phosphate depletion also increases MMP in primary and immortalized human cells and in cells of the fly midgut in vivo (Fig. 6).

Our results across the eukaryotic kingdom indicate that the higher MMP induced by phosphate deprivation contributes to improved mitochondrial energetics, morphology, and overall robustness, which could have profound implications for the many diseases, as well as the aging process itself, that are characterized by reduced MMP (Hagen et al., 1997; C. E. Hughes et al., 2020; Leprat et al., 1990; Mansell et al., 2021; Sastre et al., 1996; Sugrue & Tatton, 2001). It has been well established that phosphate limitation can extend lifespan in yeast, flies, and mice (Bergwitz et al., 2013; Ebrahimi et al., 2021; Kurosu et al., 2005, 2006). In both the fly and mouse, it was also conversely demonstrated that excessive phosphate shortens lifespan (Bergwitz et al., 2013; Kuro-o et al., 1997). In these studies, the exact mechanisms whereby phosphate abundance and sensing affect lifespan were not established. We observed increased MMP in flies treated with PFA, which limits phosphate uptake (Fig. 6E). From the evidence presented herein, as well as the previous study showing that PFA treatment extends lifespan in flies (Bergwitz et al., 2013), we hypothesize that phosphate deprivation increased MMP and overall mitochondrial health, and thereby enabled lifespan extension. This connection is supported by a recent study that demonstrated that boosting MMP by photo-activation of an ectopically expressed proton pump is sufficient to prolong lifespan in *C. elegans* (Berry et al., 2022).

This work identified genetic and environmental interventions that appear to elevate the setpoint of MMP. This improves mitochondrial protein import efficiency, results in more connected mitochondrial morphology, and reverses other deficiencies found in mutants that have low MMP. This work establishes the possibility that MMP can be modulated through cellular signaling, including augmentation of MMP above that typically observed in a wild-type cell. It also demonstrates that this MMP augmentation can occur in the absence of a functional respiratory chain. As a result, these observations lay the foundation for future studies to identify interventions that can impact the signaling mechanisms that control MMP for therapeutic benefit.

**Figure S1.**
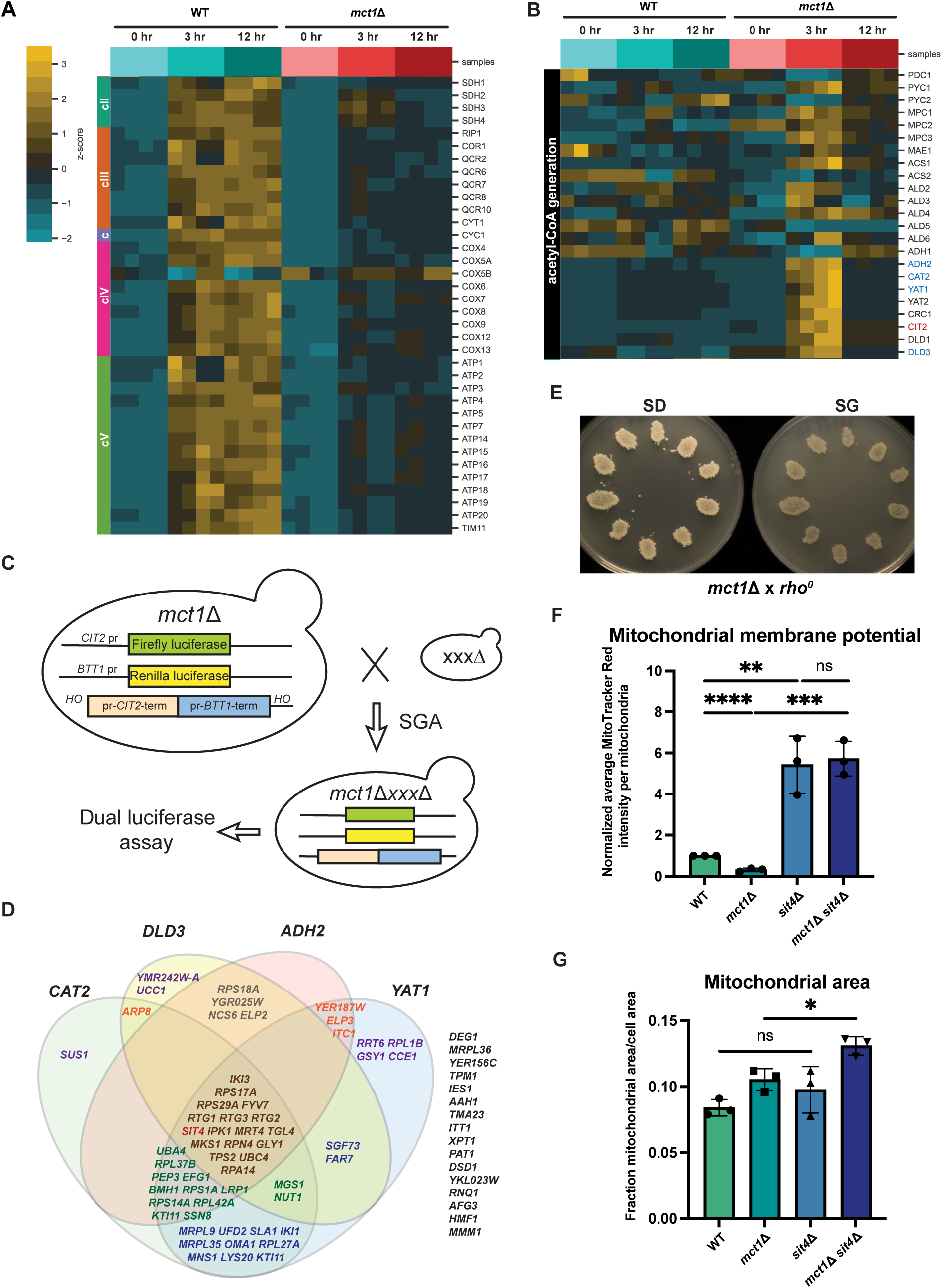
Related to Figure 1: Genetic screen to identify *SIT4* regulates nuclear responses induced in *mct1Δ* cells. (A and B) Heat map of wild-type (WT) and *mct1Δ* transcriptome. (C) Schematic of the genetic screen. The coding sequences of *CIT2* and *BTT1* were replaced with Firefly luciferase and Renilla luciferase, respectively. *CIT2* and *BTT1* genes were restored at the *HO* locus to avoid any transcriptional alterations induced by their deletion. This query strain was then crossed with a deletion library, containing ~5000 non-essential deletion strains to generate double mutants. Candidate genes were identified based on their ability to restore transcription back to wild-type levels of the Firefly:Renilla signal ratio. (D) Venn diagram of all the candidate mutants from the initial genetic screen. The inclusion of a gene in a circle indicates that in reduced expression of the indicated transcripts (*DLD3*, *CAT2*, *ADH2*, and *YAT1*) as measured by RT-qPCR. (E) Individual colonies of *mct1Δ* cells were mated with *rho^0^*cells and streaked onto a synthetic media supplemented with 2% glucose plate (SD) and then replica-plated onto a synthetic media supplemented with 2% glycerol plate (SG) to test for the presence of functional mtDNA. (F and G) Wild-type (WT), *mct1Δ*, *sit4Δ*, and *mct1Δ sit4Δ* strains were stained with MitoTracker Red and imaged. 30-90 cells were capture for each analysis. Mitochondrial membrane potential was determined by quantification of the MitoTracker Red signal that co-localized with Tom70-GFP. Mitochondrial area was calculated by the percentage of Tom70-GFP signal in total cell area. n = 3. Error bars represent the SD. Statistical significance was determined using an unpaired two-tailed t-test. ns = not significant p > 0.05; *p ≤ 0.05; **p ≤ 0.005; ***p ≤ 0.0005; ****p ≤ 0.0001.

**Figure S2.**
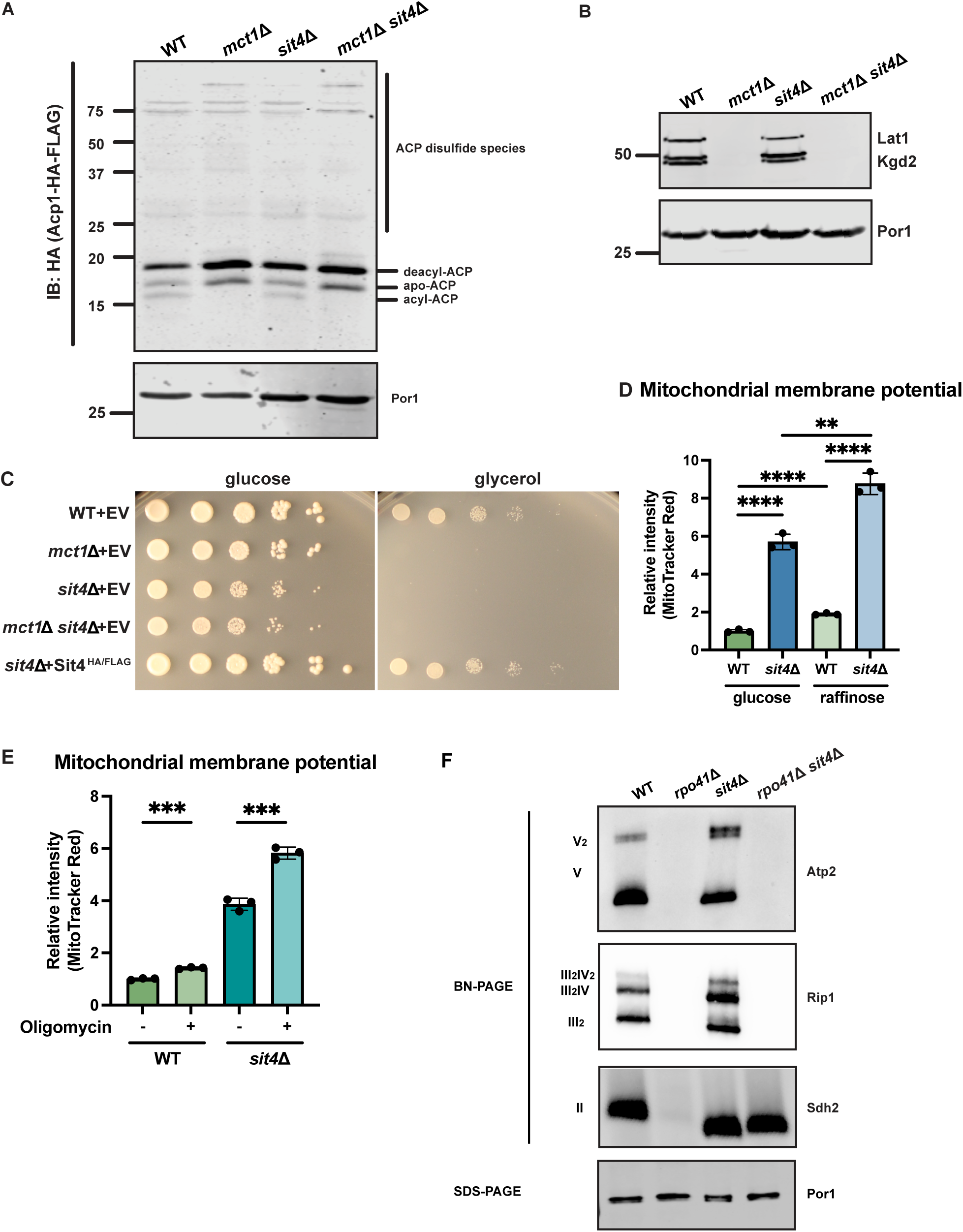
Related to Figure 2: *SIT4* deletion does not restore ACP acylation or respiratory growth in *mct1Δ* cells. (A) Mitochondria were isolated from wild-type (WT), *mct1Δ*, *sit4Δ*, and *mct1Δ sit4Δ* cells expressing Acp1-HA-FLAG from a plasmid. Acp1-HA-FLAG was immunoprecipitated with HA agarose beads and the immunoprecipitate was separated by SDS-PAGE and immunoblotted with FLAG antibody. Por1 was immunoblotted as a loading control. Original immunoblots are displayed in Figure S2—source data 1. (B) Immunoblots of isolated mitochondria from wild-type (WT), *mct1Δ*, *sit4Δ*, and *mct1Δ sit4Δ* strains blotted with antibodies against lipoic acid or porin (Por1). Two bands around the molecular weight of Kgd2 were detected. Their exact identity is unclear but could indicate different post-translation modifications of the protein. Por1 was immunoblotted as a loading control. Original immunoblots are displayed in Figure S2—source data 2. (C) Spot tests measuring the growth rate of wild-type (WT), *mct1Δ*, *sit4Δ*, and *mct1Δ sit4Δ* on either glucose- or glycerol-containing synthetic media. (D) Normalized quantification of mitochondrial membrane potential measured by flow cytometry. n = 3. Error bars represent the SD. Statistical significance was determined using an unpaired two-tailed t-test. **p ≤ 0.005; ****p ≤ 0.0001. (E) Normalized quantification of mitochondrial membrane potential measured by flow cytometry. Wild-type (WT) and *sit4Δ* cells were treated with 5 μM oligomycin for two hours. n = 3. Error bars represent the SD. Statistical significance was determined using an unpaired two-tailed t-test. ***p ≤ 0.0005. (F) Immunoblots of crude mitochondria extracted from wild-type (WT), *rpo41Δ*, *sit4Δ*, and *rpo41Δ sit4Δ* strains and separated on both BN-PAGE or SDS-PAGE. Membranes were blotted with indicated antibodies. Por1 was immunoblotted as a loading control. Original immunoblots are displayed in Figure S2—source data 3.

**Figure S3:**
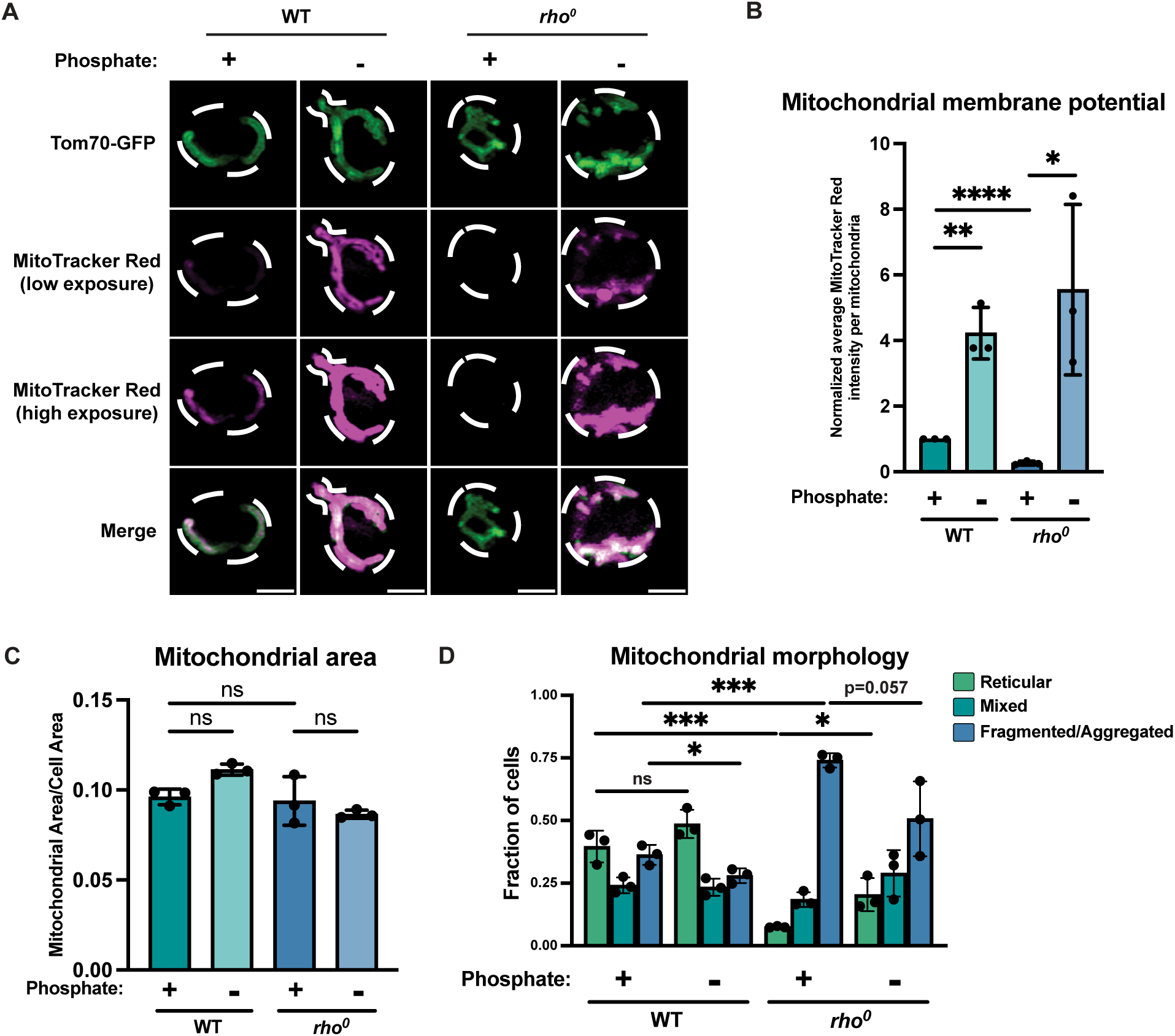
Related to Figure 3: Phosphate depletion increases mitochondrial membrane potential in wild-type and *rho^0^* cells. (A) Wild-type (WT) cells were grown in media containing high phosphate (10 mM Pi) or low phosphate (1 μM Pi) for four hours. *rho^0^* cells were grown in media containing high phosphate (+Pi, 10 mM Pi) or low phosphate (−Pi, 1 μM Pi) overnight. All cells were stained with MitoTracker Red and imaged. Representative images are shown. Scale bar represents 2 μm. (B) Quantification of (A). Mitochondrial membrane potential was determined by quantification of the MitoTracker Red signal that co-localized with Tom70-GFP. Mitochondrial area was calculated by the percentage of Tom70-GFP signal in total cell area. n = 3. Error bars represent the SD. Statistical significance was determined using an unpaired two-tailed t-test. *p ≤ 0.05; **p ≤ 0.005; ****p ≤ 0.0001. (C) Quantification of (A). Mitochondrial area was measured using Tom70-GFP signal. n = 3. Error bars represent the SD. Statistical significance was determined using a one-way ANOVA with Tukey’s multiple comparisons. ns = not significant p > 0.05. (D) Quantification of the fraction of cells imaged in (A) showing reticular, mixed, or fragmented/aggregated mitochondrial morphology. n = 3. Error bars represent the SD. Statistical significance was determined using an unpaired two-tailed t-test. ns = not significant p > 0.05; *p ≤ 0.05; ***p ≤ 0.0005.

**Figure S4:**
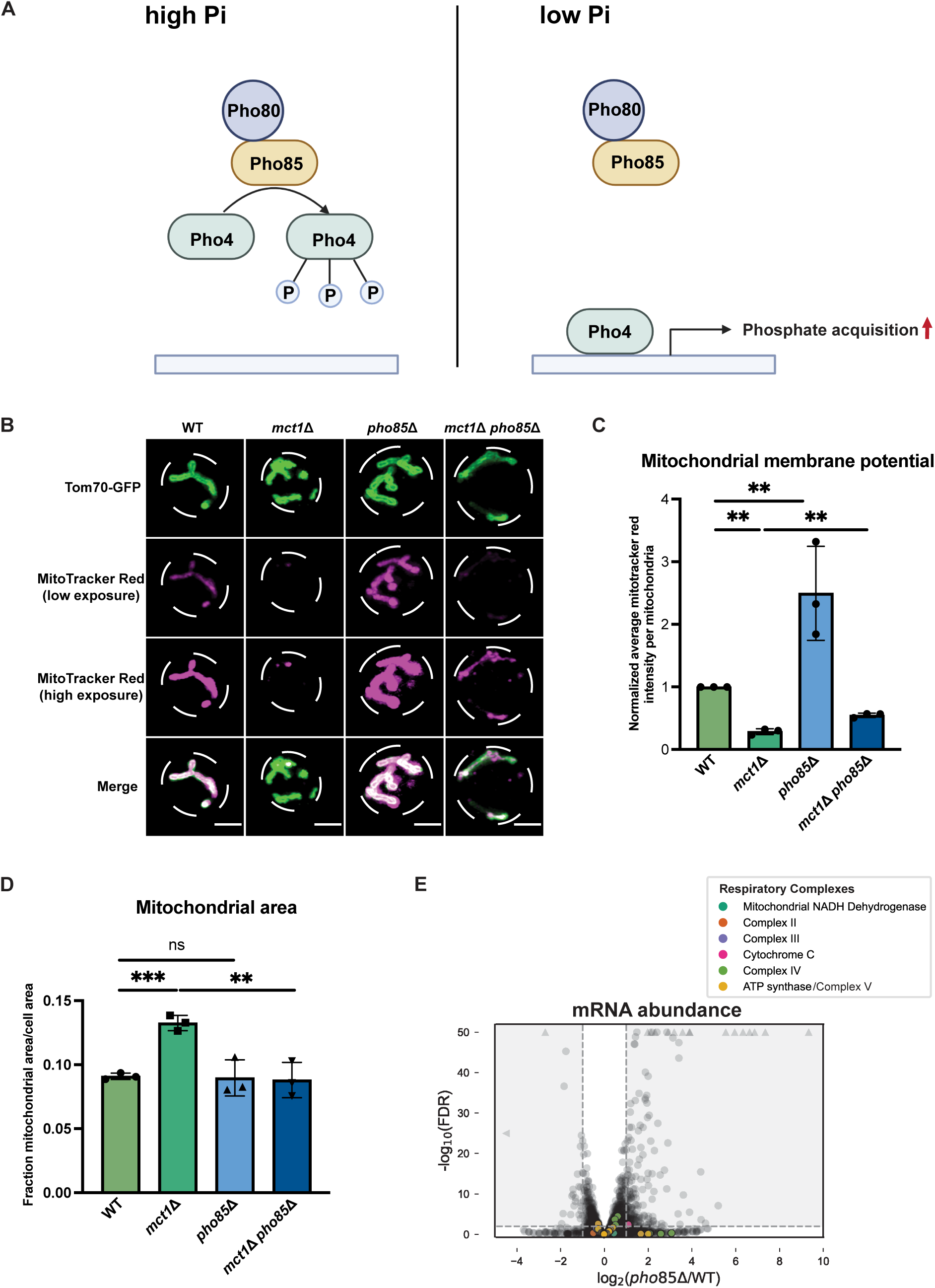
Related to Figure 5: Phosphate starvation signaling increases mitochondrial membrane potential. (A) Schematics of phosphate starvation signaling in yeast. (B) Representative images of wild-type (WT), *mct1Δ*, *pho85Δ*, and *mct1Δ pho85Δ* strains expressing Tom70-GFP from its endogenous locus stained with MitoTracker Red. Scale bar represents 2 μm. (C and D) Wild-type (WT), *mct1Δ*, *pho85Δ*, and *mct1Δ pho85Δ* strains were stained with MitoTracker Red and imaged. 30-90 cells were capture for each analysis. Mitochondrial membrane potential was determined by quantification of the MitoTracker Red signal that co-localized with Tom70-GFP. Mitochondrial area was calculated by the percentage of Tom70-GFP signal in total cell area. n = 3. Error bars represent the SD. Statistical significance was determined using an unpaired two-tailed t-test. ns = not significant p > 0.05; **p ≤ 0.005; ***p ≤ 0.0005. (E) Volcano plot of transcriptomics data of *pho85Δ* vs. WT. All components of the ETC and ATP synthase genes that were detected by RNA-seq are highlighted and color-coded.

**Figure S5.**
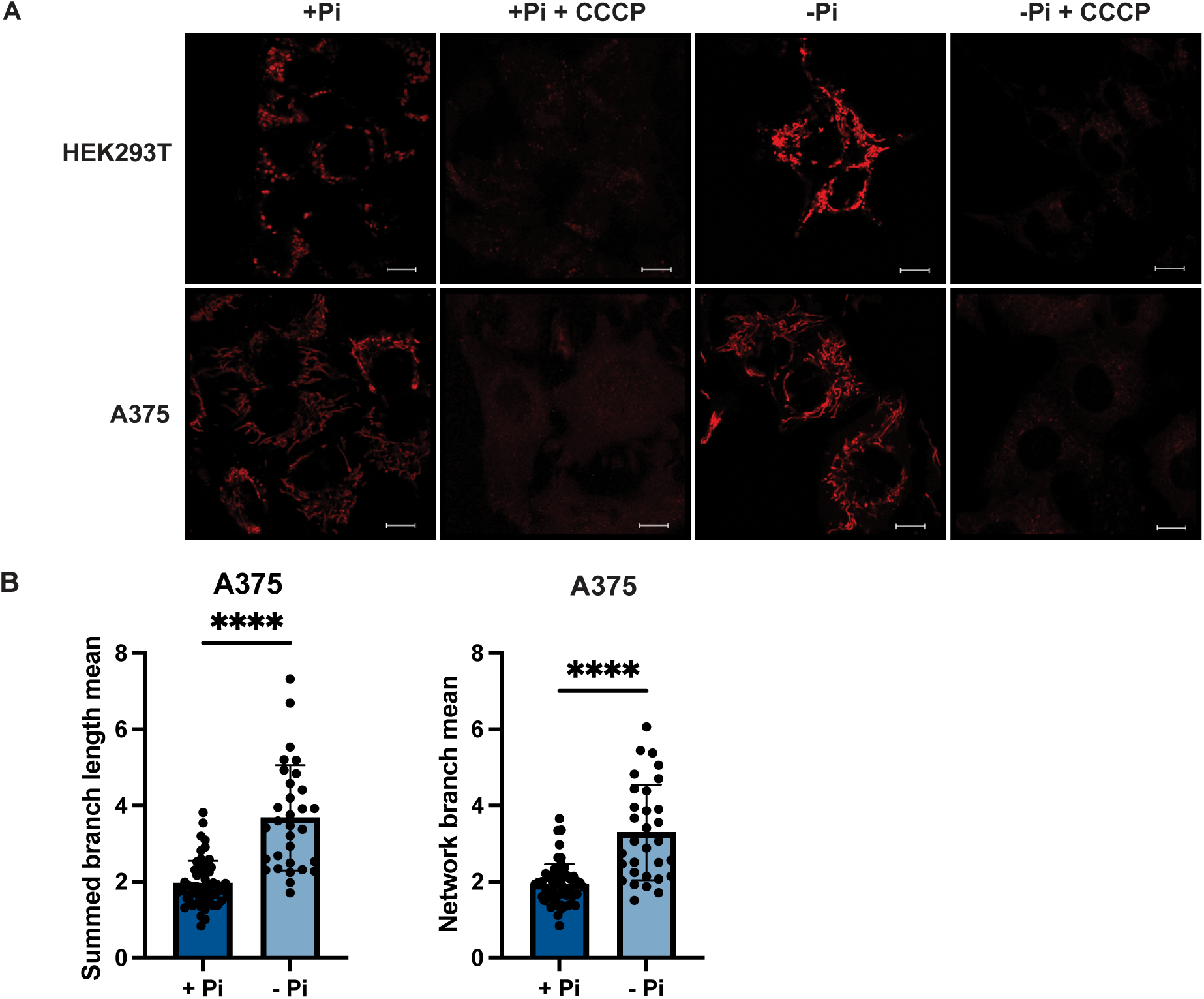
Related to Figure 6: Phosphate depletion induces elevated mitochondrial membrane potential in mammalian cells. (A) HEK293T and A375 cells were cultured in 1 mM (+Pi)- or no phosphate (−Pi)-containing media for three days. Cells were treated with 25 µM CCCP for three hours, stained with MitoTracker Red, and imaged. Representative images are shown. Scale bar represents 5 µm. (B) Summed branch lengths mean and network branch mean were measured and calculated by Mitochondrial Network Analysis (MiNA). Error bars represent the SD. Statistical significance was determined using an unpaired two-tailed t-test. ****p ≤ 0.0001.

## Methods and Materials

### Yeast strains and growth conditions

Saccharomyces cerevisiae BY4743 (MATa/a, his3/his3, leu2/leu2, ura3/ura3, met15/MET15, lys2/LYS2), was used as a parental strain to generate all knockout strains. After using PCR-based homologous recombination method, each diploid was confirmed by genotyping the targeted region in the genome. The selected diploids were dissected after 5 days sporulation at room temperature. The genotype of each haploid was determined by testing their corresponding auxotrophic or drug resistant ability. The genotypes of all strains used in this study are listed in Table I. All plasmids, antibodies, and chemicals used in this study are listed in Table II, III, and IV. Plasmids used to create yeast strains were listed in Supplementary File 1. All materials can be provided upon request.

**Table I:** Yeast strains used in this study.

| Genotype | Source | JRY identifier |
| --- | --- | --- |
| WT (BY4741) | Van Vranken et al, 2018 | JRY 2884 |
| *mct1::NatMX* | Van Vranken et al, 2018 | JRY 2885 |
| can1::STE2p-Sp_HIS5<br>lyp1del btt1::Renilla-BTT1terminator-HygMX<br>cit2::Firefly-CIT2terminator-Met15 | this study | JRY 4614 |
| ho::pr-cit2-term-pr-btt1-term-ura3 mct1::NatMX |  |  |
| can1::STE2p-Sp_HIS5<br>lyp1del btt1::Renilla-BTT1terminator-HygMX<br>cit2::Firefly-CIT2terminator-Met15 ho::pr-cit2-term-pr-btt1-term-ura3 | this study | JRY 4616 |
| *sit4*::HygMX | this study | JRY 4144 |
| *sit4*::HygMX *mct1*::NatMX | this study | JRY 4145 |
| ILV2-FLAG::KanMX | Dr. Cory Dunn | JRY 4383/CDD1084 |
| *mct1*::NatMX ILV2-FLAG::KanMX | this study | JRY 4389 |
| *sit4*::HygMX ILV2-FLAG::KanMX | this study | JRY 4385 |
| *sit4*::HygMX *mct1*::NatMX<br>ILV2-FLAG::KanMX | this study | JRY 4387 |
| *qcr2*::kanMX | this study | JRY 4556 |
| *qcr2*::kanMX *sit4*::HygMX | this study | JRY 4552 |
| *cox4*::kanMX | this study | JRY 4631 |
| *cox4*::kanMX *sit4*::HygMX | this study | JRY 4633 |
| *rpo41*::kanMX | this study | JRY 4719 |
| *rpo41::kanMX sit4::HygMX* | this study | JRY 4720 |
| *rpo41::kanMX sit4::HygMX*<br>*mct1::NatMX* | this study | JRY 4721 |
| *pho85::hisMX* | this study | JRY 4715 |
| *pho85::hisMX*<br>*mct1::NatMX* | this study | JRY 4716 |
| *pho85::hisMX* ILV2-FLAG::KanMX | this study | JRY 4743 |
| *pho85::hisMX*<br>*mct1::NatMX* ILV2-FLAG::KanMX | this study | JRY 4744 |
| *rho*<sup>0</sup> | this study | JRY 4941 |
| Tom70-yeGFP::hisMX | this study | JRY 7502 |
| Tom70-yeGFP::hisMX<br>*mct1::NatMX* | this study | JRY 7504 |
| Tom70-yeGFP::hisMX<br>*sit4::HygMX* | this study | JRY 7506 |
| Tom70-yeGFP::hisMX<br>*sit4::HygMX mct1::NatMX* | this study | JRY 7508 |
| Tom70-yeGFP::hisMX *rho*<sup>0</sup> | this study | JRY 7510 |
| Tom70-yeGFP::KanMX<br>*mct1::NatMX* | this study | JRY 7514 |
| Tom70-yeGFP::KanMX<br>*pho85::hisMX* | this study | JRY 7515 |
| Tom70-yeGFP::KanMX<br>*pho85::hisMX*<br>*mct1::NatMX* | this study | JRY 7516 |

**Table II:** Plasmids used in this study.

| Plasmids | Source |
| --- | --- |
| pRS416 Acp1-HA-FLAG | Van Vranken *et al*, 2018 |

**Table III:** Antibodies used in this study.

| Antibodies | Source |
| --- | --- |
| FLAG epitope | Sigma Aldrich, F7425 |
| Sdh2 | Dr. Dennis Winge |
| Rip1 | Dr. Dennis Winge |
| Atp2 | Dr. Dennis Winge |
| Por1 | Abcam, ab110326 |
| Lipoic acid | Abcam, ab58724 |
| Pgk1 | Abcam, ab113687 |

**Table IV:**
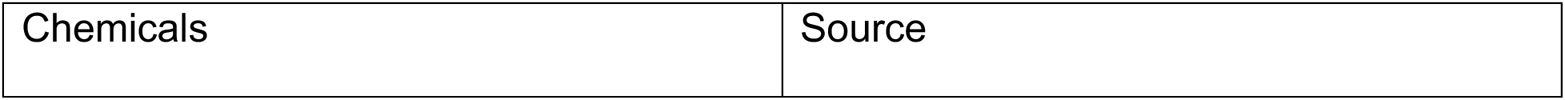
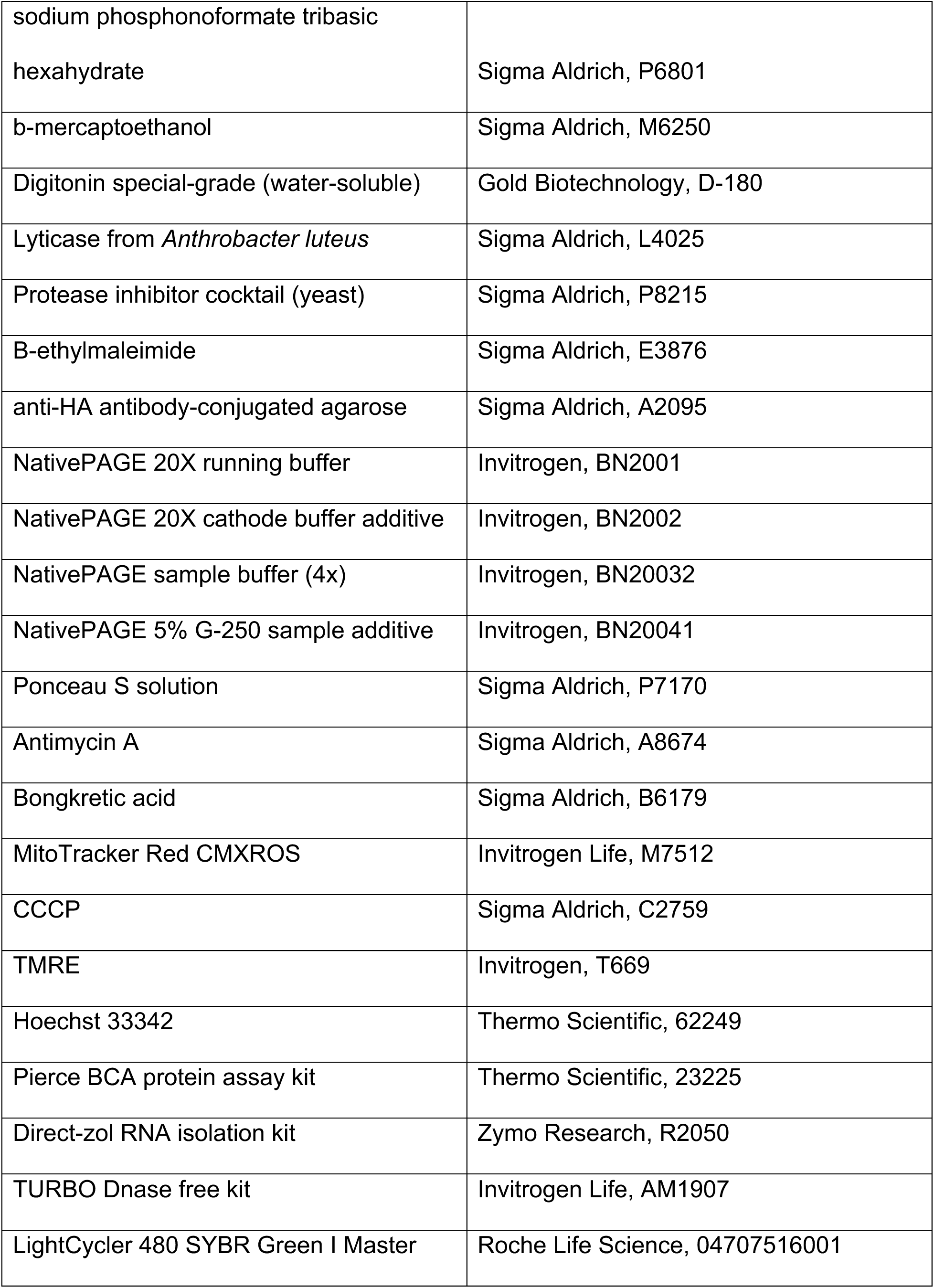
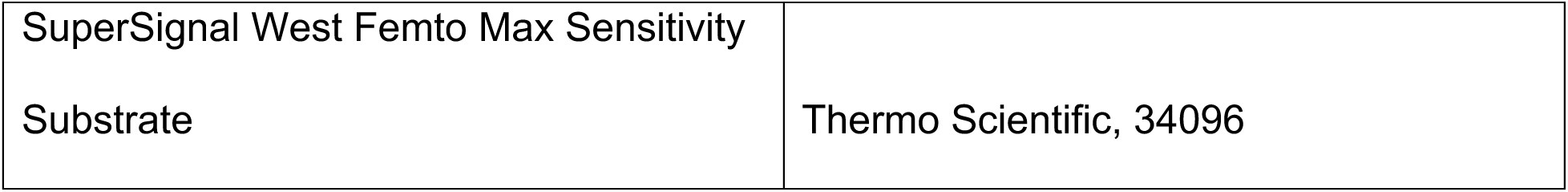
Chemicals and commercial kits used in this study.

Yeast transformation was performed by the LiAc/TE method, and successfully transformed cells were selected on agar plates containing synthetic complete media with 2% glucose (SD) lacking the corresponding amino acid(s).

*rho^0^* cells were generated using 25 ug/ml of ethidium bromide (Sigma, E7637). Small colonies were picked to be validated with the following two assays. The *rho^0^* cells failed to grow on glycerol containing media. The isolated DNA from *rho^0^* cells yielded no product after 45 cycles of PCR with primers amplifying small fragments of *COX2* (forward: 5’-GAATGATGTACCAACACCTTATG-3’, reverse: 5’-GATACTGCTTCGATCTTAATTGGC-3’) and *ATP6* (forward: 5’-GACTATTATTTGGTTTACAATCATC-3’, reverse: 5’-TTAATGTAAGTATACTGCATCTTTTAAATATG-3’), while the DNA from wild-type cells generated robust PCR product.

For most assays, unless specified, yeast cells were grown in synthetic media with 2% glucose overnight and backdiluted to a much lower OD (the exact number of cells dependent on the assays and mutants being used) in media indicated in each assay. For assays that used carbon sources other than glucose, the carbon source and percentage was specified in the figure or figure legend. Cells were harvested between 0.2 - 0.6 OD to ensure similar metabolic state. For phosphate depletion assay, saturated or active growing yeast cells (below 0.6 OD) were washed twice with a much larger volume of water. The cells were then grown in synthetic media without inorganic phosphate supplemented with KCl (Formedium, CYN6701) with amino acids, 2% glucose, and the indicated amount of potassium phosphate monobasic, pH 4.1.

### Mammalian cells and growth conditions

HEK293T and A375 cell lines were cultured and maintained in Dulbecco’s modified Eagle medium (DMEM) supplemented with 10% FBS in an incubator at 37 °C with 5% CO_2_.

To deplete phosphate, cells were washed at minimum three times with filter-sterilized normal saline (0.9% NaCl) and trypsinized with 0.25% trypsin in citrate saline (STEMCELL Technologies, 07400). Cells were resuspended with DMEM with no phosphate (Thermofisher Scientific, 11971025), supplemented with 10% One Shot Dialyzed FBS (Thermofisher Scientific, A3382001) and 2 mM sodium pyruvate, hereby known as −Pi media, divided equally by cell number into 15 mL falcon tubes, and centrifuged to pellet cells.

Cells were resuspended in either the aforementioned −Pi media, or in +Pi media, which is media supplemented with 1 mM sodium phosphate monobasic, hereby known as +Pi media, depending on condition cells were to be plated in. Cells were grown in either −Pi or +Pi media in an incubator at 37 °C and 5% CO_2_ for three days prior to collection.

### Fly stocks and growth condition

*Drosphila melanogaster* stain *W^1118^* (BDSC 3605) was used for testing the effect of phosphate uptake on mitochondrial membrane potential. The flies were maintained on fly semi-defined food which consists of 1% Agar, 8% Yeast, 3% Sucrose, 6% Glucose, 0.05% MgSO4, 0.05% CaCl_2_, 1% Tegosept, and 0.6% propionic acid.

### Yeast genetic screen library construction and dual luciferase assay

The deletion collection (haploid) was mated with the query strain (*can1*::STE2p-Sp_HIS5 lyp1del *btt1*::Rluc-BTT1terminator-HygMX *cit2*::Fluc-CIT2terminator-Met15 *ho*:: pr-cit2-term-pr-btt1-term-ura3 *mct1*::NatMX) on YPAD plates and selected for diploid on YPAD + G418/Nat/Hyg for 2 days at 30 °C. Selected diploids were sporulated on enriched sporulation media (20 g/l agar, 10 g/l potassium acetate, 1 g/l yeast extract, 0.5 g/l glucose, 0.1 g/l amino-acids supplement) for 5 days at room temperature.

Desired haploids were selected on SD-his/arg/lys + canavanine/thialysine for 2 days at 30 °C followed by growing on SD without ammonium sulfate supplemented with monosodium glutamic acid (MSG)-his/arg/lys + canavanine/thialysine/G418 for 1 day at 30 °C. The final selection was conducted on SD/MSG-his/arg/lys/ura/cys/met + canavanine/thialysine/G418/Nat/Hyg for 1 day at 30 °C.

Cells were grown in 384-well plate overnight in SD complete (2% glucose). After back-diluting to around OD 0.1, the cells were grown for 6 hours in SR complete media (2% raffinose). The dual luciferase assay was conducted using the Dual-Glo Luciferase Assay System (Promega) following the product manual. Both firefly and Renilla luciferase was measured using a GloMax plate reader (Promega) with an injector in 96-well plate format.

### Mitochondrial membrane potential measurement in yeast

0.2 OD of yeast cells were pelleted down and incubated in the same growth media containing 100 nM of MitoTracker Red CMXRos (Life Technologies) for 30 min at room temperature. Cells were spun down again and resuspended in the same media, which was either imaged by fluorescence microscopy or measured by fluorescence-assisted cell sorting. All experiments were performed with three biological triplicates.

### MitoTracker Red staining in mammalian cells

Cells were washed 3x and trypsinized as previously stated, and then resuspended in −Pi or +Pi media with a 20 nM final concentration of MitoTracker Red, then incubated at 37 °C for 15 minutes. Cells are centrifuged, washed once with normal saline, centrifuged, and resuspended in −Pi or +Pi media and proceed with flow cytometry analysis. All experiments were performed with three biological triplicates.

### MitoTracker Red staining and fluorescence microscopy in rat primary hepatocytes

Rat Primary hepatocytes (Wister, Lonza, RICP01) were cultured as instructed. In brief, 0.9 million primary hepatocytes were plated in collagen coated florodish in HCM^TM^ SingleQuots^TM^ Kit (Lonza, CC-4182, containing ascorbic acid, bovine serum albumin-fatty acid free (BSA-FAF), hydrocorticosone, human epidermal growth factor (hEGF), transferrin, insulin and gentamicin/amphotericin-B (GA)), overnight at 37 °C. Cells were treated with 3 mM or 1 mM PFA in DMEM with 10% FBS and 1% P/S for 24 hours. Cells were then washed two times with saline-0.9% NaCl before staining with 20 nM MitoTracker Red CMXRos at 37 °C for 15 minutes. After two saline washes, live imaging of MitoTracker Red fluorescence was done in DMEM with 10% FBS on Zeiss 900 Airyscan. The intensity of MitoTracker Red for 80 cells per treatment was quantified using Fiji. Cellular boundaries were defined using the Free hand tool, and background intensity was subtracted. The graphs were plotted using Prism v9.

### TMRE staining and fluorescence microscopy in adult *Drosophila* gut intestines

3-day old flies (15 females and 10 males) were transferred in vials with semi-defined media with or without 1 mM sodium phosphonoformate tribasic hexahydrate (PFA) (Sigma, P6801). After 2 weeks, fly guts (midgut, R4-R5 section) from both control and PFA-treated group were dissected in Shields and Sang M3 Insect Media (Sigma, S8398) with or without 1 mM PFA added. Dissected guts were staining with 1 μM TMRE (Sigma, 87917) in the same media condition for 20 minutes at room temperature. After 2 washes with the corresponding media supplemented with 1 μM Hoechst 33342 (Thermo Scientific, 62249) and 0.01 μM TMRE, live imaging of gut was performed on Zeiss 900 Airyscan. Intensity of TMRE was quantified in Fiji, and background was subtracted. Figures was prepared in Prism v9.

### Fluorescence microscopy

Yeast containing Tom70-GFP tagged chromosomally at its endogenous locus were grown and stained with MitoTracker Red CMXRos as described for flow cytometry. In short, 5 × 10^6^ yeast were harvested by centrifugation and resuspended in 1 ml media containing 0.2 μM MitoTracker Red CMXRos. Cells were incubated in the dark at room temperature for 30 minutes, washed once in 1 ml media, and resuspended in 20 μl of media.

For quantitative microscopy, images were collected on an Axio Observer (Carl Zeiss) with a 63x oil-immersion objective (Carl Zeiss, Plan Apochromat, NA 1.4) and an Axiocam 503 mono camera (Carl Zeiss). Three optical z-sections across 1 μm sections were collected per image. Each image contained on average 19 yeast cells (range: 5 to 49 yeast cells). Quantifications were derived from average values per picture. Five pictures were averaged for each condition in an experimental replicate. Final values are averages of three experimental replicates. Images were collected in ZEN (Carl Zeiss) and processed in Fiji (Schindelin et al., 2012). All quantifications were done on maximum-intensity projections. All images within each experiment were processed identically.

For mitochondrial membrane potential quantifications, a mask was created from thresholded Tom70-GFP images and used to measure the average MitoTracker Red CMXRos fluorescence intensity of each mitochondria. Each condition within an experimental replicate was normalized to untreated wild type yeast to control for variations in MitoTracker Red CMXRos staining.

For mitochondrial mass quantifications, mitochondrial area per picture was measured from thresholded Tom70-GFP images. Cell area per picture was measured from Tom70-GFP images at a threshold such that the entirety of each cell was outlined. Plots are mitochondrial area divided by cell area.

For mitochondrial morphology quantifications, the contrast of Tom70-GFP images were adjusted for optimal viewing and morphology scored by an unblinded researcher.

For image panels, super-resolution Airyscan images were collected using an LSM800 (Carl Zeiss) equipped with an Airyscan detector and a 63x oil-immersion objective (Carl Zeiss, Plan Apochromat, NA 1.4). Optical z-sections were acquired across the entire yeast cell at a step size optimal for Airyscan super-resolution, 0.15 μm. Images were acquired on ZEN software and processed using the automated Airyscan processing algorithm in ZEN (Carl Zeiss). Maximum-intensity projections and contrast enhancement were done in Fiji. Contrast changes were kept identical within an experiment.

HEK 293T and A375 cells were seeded on 35 mm Fluorodish plates (World Precision Instruments). 24 hr later, cells were stained with 20 nM final concentration of MitoTracker Red CMXRos for 20 minutes, washed once in PBS, and were live-cell imaged on a Zeiss 880 confocal laser scanning microscope incubated at 37 °C, 5% CO_2_. All mammalian cell imaging was performed with a Plan-Apochromat 63x/1.40 Oil DIC f/ELYR objective. Images were Airyscan processed using the Zeiss Zen Blue software.

### Fluorescence-assisted cell sorting

Cells were stained with MitoTracker Red CMXRos. A total of 10,000 events were measured on a BD FACSCanto with BD FACSDiva 8.0.1.1 (BD Biosciences). The median fluorescence values were plotted with Prism 9.

### Mitochondrial morphology quantification using MiNA

The MiNA plugin (Valente et al., 2017) was installed in Fiji. MitoTracker Red signaling of each cell was outlined as ROI and the skeletonized mitochondria were generated. Then, the mean summed branch lengths and the mean network branch was automatically measured and calculated. Around 30-60 cells were analyzed for each condition. The final bar graphs were plotted with Prism 9.

### Mitochondrial protein import assay

10 OD of total culture were harvested at an OD_600_ between 0.3 and 0.5. Cell pellets were washed with water and lysed in 500 μl of 2 M lithium acetate for 10 minutes on ice. Lysed cells were resuspended in 500 μl of ice cold 0.4 M NaOH and left on ice for 10 minutes. The pellets were resuspended with 250 μl of 2x Laemmeli buffer with 5% BME. The lysate was boiled for 5 minutes, and the supernatant was loaded on a 12% SDS-PAGE gel and assessed by immunoblot.

### Western blotting

Whole cell or mitochondria extract were separated on an SDS-PAGE or BN-PAGE gel and transferred to nitrocellulose membranes (SDS-PAGE) or activated PVDF membranes (BN-PAGE) with a Power Station (Biorad). Membranes were blocked in blocking buffer (Tris buffered saline (50 mM Tris-HCl pH 7.4, 150 mM NaCl, 5% non-fat dry milk) and probed with the primary antibodies listed in Table III and the secondary antibodies. Antibodies were either visualized with Licor Odyssey or SuperSignal Enhanced Chemiluminescence Solution (Thermo Scientific, 34096) and a Chemidoc MP System (BioRad).

### Crude mitochondrial isolation

Crude mitochondrial isolation was performed as described previously (Van Vranken et al., 2018). Cell pellets were resuspended in TD buffer (100 mM Tris-SO4, pH 9.4 and 100 mM DTT) and incubated for 15 min at 30 °C. Cells were then washed once in SP buffer (1.2 M Sorbitol and 20 mM potassium phosphate, pH 7.4) and incubated in SP buffer with 0.3 mg/mL lyticase (Sigma, L4025) for 1 hr at 30 °C to digest the cell wall. Spheroplasts were washed once and homogenized in ice-cold SEH buffer (0.6 M sorbitol, 20 mM HEPES-KOH, pH 7.4, 1 mM PMSF, yeast protease inhibitor cocktail (Sigma, P8215)) by applying 20 strokes in a dounce homogenizer. Crude mitochondria were isolated by differential centrifugation at 3000 xg first to remove larger debris and 10,000 xg to pellet down mitochondria. Protein concentrations were determined using a Peirce BCA Protein Assay Kit (Thermo Scientific, 23225).

### Blue native polyacrylamide gel electrophoresis (BN-PAGE)

BN-PAGE was performed as described previously (Van Vranken et al., 2018). 100 μg of mitochondria were resuspended in 1x lysis buffer (Invitrogen, BN20032) supplemented with yeast protease inhibitor cocktail (Sigma, P8215) and solubilized with 1% digitonin for 20 min on ice. Solubilized mitochondria were cleared by centrifugation at 20,000 xg for 20 min. Lysate were mixed with NativePAGE 5% G-250 Sample Additive (Invitrogen, BN20041) and resolved on a 3%-12% gradient native gel (Invitrogen, BN1001BOX).

### Mitochondrial isolation and immunoprecipitation for the ACP acylation study

Mitochondrial isolation and immunoprecipitation were performed as described previously (Van Vranken et al., 2018). Briefly, cell pellets were washed and lysed as described in the “Crude mitochondrial isolation” section of this manuscript. Spheroplasts were washed once and homogenized in ice-cold SEH buffer (0.6 M sorbitol, 20 mM HEPES-KOH, pH 7.4, 1 mM PMSF, yPIC) with 10 mM N-ethylmaleimide (NEM) (Sigma, E3876) by applying 20 strokes in a dounce homogenizer. Crude mitochondria were isolated by differential centrifugation. Protein concentrations were determined using a Peirce BCA Protein Assay Kit (Thermo Scientific). 1 mg of crude mitochondria were resuspended in 200 μl of XWA buffer (20 mM HEPES, 10 mM KCl, 1.5 mM MgCl_2_, 1 mM EDTA, 1 mM EGTA, pH 7.4) with 10 mM NEM and 0.7% digitonin added and incubated on ice for 30 min. After centrifugation at 20,000 xg for 20 mins, solubilized mitochondria were incubated with pre-equilibrated anti-HA antibody-conjugated agarose (Sigma, A2095) for 2 hr at 4 °C. The agarose was washed 3 times and eluted by incubating in 2x Laemmeli buffer at 65 °C for 10 min. Elution was isolated by SDS-PAGE and subjected for immunoblot analysis.

### RNA isolation and qPCR

RNA was isolated as described previously (Zurita Rendón et al., 2018, p. 1). Briefly, cell pellets were resuspended in Trizol reagent (Ambion, 15596026) and bead bashed to lyse the cell (20 s bash with 30 s break on ice, 6 cycles). Equal volume of ethanol was added to the sample and RNA is isolated using the Direct-zol kit (Zymo research, R2050). The RNA eluted from the column was treated with TURBO DNase kit (Invitrogen, AM1907) to remove remaining DNA contamination. After normalization of RNA content, cDNA was generated by using a High-capacity cDNA Reverse Transcription kit (Applied Biosystem, 4368813). Quantative PCR was performed using the LightCycler 480 SYRB Green I Master (Roche, 04707516001). The raw data was analyzed by absolute quantification/2^nd^ derivative of three independent biological replicates with each being the average of three technical replicates.

### RNA-sequencing

For dataset GSE151606, yeast cultures were initially grown in synthetic media supplemented with 2% glucose, then removed from original media and transferred to synthetic media with 2% raffinose. Cultures were flash frozen and later total RNA was isolated using the Direct-zol kit (Zymo Research, R2050) with on-column DNase digestion and water elution. Sequencing libraries were prepared by purifying intact poly(A) RNA from total RNA samples (100-500 ng) with oligo(dT) magnetic beads and stranded mRNA sequencing libraries were prepared as described using the Illumina TruSeq Stranded mRNA Library Preparation Kit (RS-122-2101, RS-122-2102). Sequencing libraries (25 pM) were chemically denatured and applied to an Illumina HiSeq v4 single read flow cell using an Illumina cBot. Hybridized molecules were clonally amplified and annealed to sequencing primers with reagents from an Illumina HiSeq SR Cluster Kit v4-cBot (GD-401-4001). 50 cycle single-read sequence run performed using HiSeq SBS Kit v4 sequencing reagents (FC-401-4002). Read pre-processing was performed by Fastp, v0.20.0 (S. Chen et al., 2018). Read alignment was performed by STAR, v2.7.3a (Dobin et al., 2013). Read quantification was performed by htseq, v0.11.3 (Anders et al., 2015). Read QC was performed by fastqc, v0.11.9 (Andrews S., 2010). Total QC was performed by multiqc, v1.8 (Andrews S., 2010). Library complexity QC was performed by dupradar, v1.10.0 (Sayols et al., 2016). Genome_build Ensembl R64-1-1 (GCA_000146045.2), version 100 was used during alignment and quantification. Scripts for data processing can be found at https://github.com/j-berg/ouyang_analysis_2022/tree/main/rnaseq/GSE151606_mct1_timecourse.

For dataset GSE209726, yeast cultures were grown in SD-complete overnight and harvested at OD_600_ between 0.2 - 0.4. Intact poly(A) RNA was purified from total RNA samples (100-500 ng) with oligo(dT) magnetic beads. Stranded mRNA sequencing libraries were prepared as described using the Illumina TruSeq Stranded mRNA Library Prep kit (20020595) and TruSeq RNA UD Indexes (20022371). Purified libraries were qualified on an Agilent Technologies 2200 TapeStation using a D1000 ScreenTape assay (cat# 5067-5582 and 5067-5583). The molarity of adapter-modified molecules was defined by quantitative PCR using the Kapa Biosystems Kapa Library Quant Kit (cat#KK4824). Individual libraries were normalized to 1.30 nM in preparation for Illumina sequence analysis. Sequencing libraries were chemically denatured and applied to an Illumina NovaSeq flow cell using the NovaSeq XP workflow (20043131). Following transfer of the flowcell to an Illumina NovaSeq 6000 instrument, a 150 × 150 cycle paired end sequence run was performed using a NovaSeq 6000 S4 reagent Kit v1.5 (20028312). Read pre-processing was performed by Fastp, v0.20.1(S. Chen et al., 2018). Read alignment was performed by STAR, v2.7.7a (Dobin et al., 2013). Read postprocessing was performed by samtools v1.11 (H. Li et al., 2009). Read quantification was performed by htseq, v0.13.5 (Anders et al., 2015). Genome_build Ensembl R64-1-1 (GCA_000146045.2), version 100 was used during alignment and quantification. Scripts for data processing can be found at https://github.com/j-berg/ouyang_analysis_2022/tree/main/rnaseq/GSE209726_mct1_sit4_deletions.

For dataset GSE212790, yeast were grown in the indicated media overnight and harvested between OD_600_=0.2-0.4, with a total OD of 5 per sample. After QC procedures, mRNA from eukaryotic organisms is enriched from total RNA using oligo(dT) beads. The mRNA is then fragmented randomly in fragmentation buffer, followed by cDNA synthesis using random hexamers and reverse transcriptase. After first-strand synthesis, a custom second-strand synthesis buffer (Illumina) is added, with dNTPs, RNase H and Escherichia coli polymerase I to generate the second strand by nick-translation and AMPure XP beads is used to purify the cDNA. The final cDNA library is ready after a round of purification, terminal repair, Atailing, ligation of sequencing adapters, size selection and PCR enrichment. Library concentration was first quantified using a Qubit 2.0 fluorometer (Life Technologies), and then diluted to I ng/gl before checking insert size on an Agilent 2100 and quantifying to greater accuracy by quantitative PCR (Q-PCR) (library activity >2 nM). Libraries are fed into Novaseq6000 machines according to activity and expected data volume. A paired-end 150 bp sequencing strategy was used and all samples were sequenced to at least 6 Gb. XPRESSpipe v0.6.3 (Berg et al., 2020) was used to process sequence files, with the following command: xpresspipe peRNAseq … -a AGATCGGAAGAGCGTCGTGTAGGGAAAGAGTGTAGATCTCGGTGGTCGCCGTATC ATT GATCGGAAGAGCACACGTCTGAACTCCAGTCACGGATGACTATCTCGTATGCCGT CTTCTGCTTG --sjdbOverhang 149 --quantification_method htseq --remove_rrna. Genome_build Ensembl R64-1-1 (GCA_000146045.2), version 106 was used during alignment and quantification. Scripts for data processing can be found at https://github.com/j-berg/ouyang_analysis_2022/tree/main/rnaseq/GSE212790_genetic_nutrient_perturbation.

### Data analysis and statistics for RNA-sequencing

Analysis code notebooks can be accessed at https://github.com/j-berg/ouyang_analysis_2022. Differential expression analysis was performed using DESeq2 (Love et al., 2014) with the FDR threshold (α) set at 0.1. Data visualization was performed in Python using Pandas (McKinney, 2010), numpy (Oliphant, 2006; van der Walt et al., 2011), scikit-learn (Buitinck et al., 2013), matplotlib (Hunter, 2007), seaborn (Waskom et al., 2012).

### Sample Preparation for Mass Spectrometry

Yeast proteomes were extracted using a buffer containing 200 mM EPPS, 8 M urea, 0.1% SDS, and 1X protease inhibitor [Peirce protease inhibitor mini tablets]). 100 μg of each proteome was prepared as follows. 10 mM tris(2-carboxyethyl)phosphine hydrochloride) and incubated at room temperature for 10 mins. Iodoacetimide was added to a final concentration of 10 mM to each sample and incubated for 25 mins. in the dark. Finally, DTT was added to each sample to a final concentration of 10 mM. A buffer exchange was carried out using a modified SP3 protocol (C. S. Hughes et al., 2014, 2019). Briefly, ~500 μg of each SpeedBead Magnetic Carboxylate modified particles (Cytiva; 45152105050250, 65152105050250) mixed at a 1:1 ratio was added to each sample. 100% ethanol was added to each sample to achieve a final ethanol concentration of at least 50%. Samples were incubated with gentle shaking for 15 mins. Samples were washed three times with 80% ethanol. Protein was eluted from SP3 beads using 200 mM EPPS pH 8.5 containing trypsin (ThermoFisher Scientific) and Lys-C (Wako). Samples were digested overnight at 37°C with vigorous shaking. Acetonitrile was added to each sample to achieve a final concentration of 30%. Each sample was labelled, in the presence of SP3 beads, with ~250 μg of TMTpro-16plex reagents (ThermoFisher Scientific) (J. Li et al., 2020; Thompson et al., 2019) for 1 hour. Following confirmation of satisfactory labelling (>97%), excess TMTpro reagents were quenched by addition of hydroxylamine to a final concentration of 0.3%. The full volume from each sample was pooled and acetonitrile was removed by vacuum centrifugation for one hour. The pooled sample was acidified using formic acid and peptides were de-salted using a Sep-Pak Vac 200 mg tC18 cartridge (Waters). Peptides were eluted in 70% acetonitrile, 1% formic acid and dried by vacuum centrifugation. Phosphopeptides were enriched using a Hugh Select Phosphopeptide Enrichment Kit (ThermoFisher Scientific). Flow through from the phosphopeptide enrichment column was collected for whole proteome analysis. The peptides were resuspended in 10 mM ammonium bicarbonate pH 8, 5% acetonitrile and fractionated by basic pH reverse phase HPLC. In total 24 fractions were collected. The fractions were dried in a vacuum centrifuge, resuspended in 5% acetonitrile, 1% formic acid and desalted by stage-tip. Final peptides were eluted in, 70% acetonitrile, 1% formic acid, dried, and finally resuspended in 5% acetonitrile, 5% formic acid. In the end, 8 fractions were analyzed by LC-MS/MS.

### Mass spectrometry data acquisition

Data were collected on an Orbitrap Eclipse mass spectrometer (ThermoFisher Scientific) coupled to a Proxeon EASY-nLC 1000 LC pump (ThermoFisher Scientific). Whole proteome peptides were separated using a 90-min gradient at 500 nL/min on a 30-cm column (i.d. 100 μm, Accucore, 2.6 μm, 150 Å) packed in house. High-field asymmetric-waveform ion mobility spectroscopy (FAIMS) was enabled during data acquisition with compensation voltages (CVs) set as −40 V, −60 V, and −80 V (Schweppe et al., 2019). MS1 data were collected using the Orbitrap (60,000 resolution; maximum injection time 50 ms; AGC 4 × 10^5^). Determined charge states between 2 and 6 were required for sequencing, and a 60 s dynamic exclusion window was used. Data dependent mode was set as cycle time (1 s). MS2 scans were performed in the Orbitrap with HCD fragmentation (isolation window 0.5 Da; 50,000 resolution; NCE 36%; maximum injection time 86 ms; AGC 1 × 10^5^). Phosphopeptides were separated using a 120 min gradient at 500 nL/min on a 30-cm column (i.d. 100 μm, Accucore, 2.6 μm, 150 Å) packed in house. The phosphopeptide enrichment was injected twice using two different FAIMS methods. For the first injection, the FAIMS CVs were set to −45 V and −65 V. For the second injection, the FAIMS CVs were set to −40 V, −60 V, and −80 V (Schweppe et al., 2019). For both methods, MS1 data were collected using the Orbitrap (120,000 resolution; maximum ion injection time 50 ms, AGC 4×10^5^). Determined charge states between 2 and 6 were required for sequencing, and a 60 s dynamic exclusion window was used. Data dependent mode was set as cycle time (1 s). MS2 scans were performed in the Orbitrap with HCD fragmentation (isolation window 0.5 Da; 50,000 resolution; NCE 36%; maximum injection time 250 ms; AGC 1 × 10^5^).

### Phosphoproteomics data analysis

Raw files were first converted to mzXML, and monoisotopic peaks were re-assigned using Monocle (Rad et al., 2021). Searches were performed using the Comet search algorithm against a yeast database downloaded from Uniprot in June 2014. We used a 50 ppm precursor ion tolerance and 0.9 Da product ion tolerance for MS2 scans collected in the ion trap and 0.02 Da product ion tolerance for MS2 scans collected in the Orbitrap. TMTpro on lysine residues and peptide N-termini (+304.2071 Da) and carbamidomethylation of cysteine residues (+57.0215 Da) were set as static modifications, while oxidation of methionine residues (+15.9949 Da) was set as a variable modification. For phosphorylated peptide analysis, +79.9663 Da was set as a variable modification on serine, threonine, and tyrosine residues.

Peptide-spectrum matches (PSMs) were adjusted to a 1% false discovery rate (FDR) (Elias & Gygi, 2007). PSM filtering was performed using linear discriminant analysis (LDA) as described previously (Huttlin et al., 2010), while considering the following parameters: comet log expect, different sequence delta comet log expect (percent difference between the first hit and the next hit with a different peptide sequence), missed cleavages, peptide length, charge state, precursor mass accuracy, and fraction of ions matched. Each run was filtered separately. Protein-level FDR was subsequently estimated at a data set level. For each protein across all samples, the posterior probabilities reported by the LDA model for each peptide were multiplied to give a protein-level probability estimate. Using the Picked FDR method (Savitski et al., 2015), proteins were filtered to the target 1% FDR level. Phosphorylation site localization was determined using the AScore algorithm (Beausoleil et al., 2006).

For reporter ion quantification, a 0.003 Da window around the theoretical *m/z* of each reporter ion was scanned, and the most intense *m/z* was used. Reporter ion intensities were adjusted to correct for the isotopic impurities of the different TMTpro reagents according to manufacturer specifications. Peptides were filtered to include only those with a summed signal-to-noise (SN) of 160 or greater across all channels. For each protein, the filtered peptide TMTpro SN values were summed to generate protein quantification.

### Data availability

The mass spectrometry data have been deposited to the ProteomeXchange Consortium with the data set identifier PXD037405. RNA sequencing data have been deposited to the GEO Omnibus Repository with data set identifiers GSE151606, GSE212790, and GSE209726. Code for high-throughput dataset analysis is archived on GitHub at https://github.com/j-berg/ouyang_analysis_2022, and at Zenodo at https://doi.org/10.5281/zenodo.7212729.

## Acknowledgements

We thank University of Utah core facilities, especially James Marvin, PhD at the Flow Cytometry Core, Brian Dalley, PhD at the High-Throughput Genomics Core, and the DNA/Peptide Synthesis Core. We thank members of the Rutter lab for discussion and feedback on the manuscript. Several of the figures were created with BioRender.com.

## Reference

Abdel-Fattah, W., Jablonowski, D., Santo, R. D., Thüring, K. L., Scheidt, V., Hammermeister, A., Have, S. ten, Helm, M., Schaffrath, R., & Stark, M. J. R. (2015). Phosphorylation of Elp1 by Hrr25 Is Required for Elongator-Dependent tRNA Modification in Yeast. PLOS Genetics, 11(1), e1004931. https://doi.org/10.1371/journal.pgen.1004931

Anders, S., Pyl, P. T., & Huber, W. (2015). HTSeq—A Python framework to work with high-throughput sequencing data. Bioinformatics, 31(2), 166–169. https://doi.org/10.1093/bioinformatics/btu638

Andrews S. FastWC; 2010. https://www.bioinformatics.babraham.ac.uk/projects/fastqc/

Angerer, H., Schönborn, S., Gorka, J., Bahr, U., Karas, M., Wittig, I., Heidler, J., Hoffmann, J., Morgner, N., & Zickermann, V. (2017). Acyl modification and binding of mitochondrial ACP to multiprotein complexes. Biochimica et Biophysica Acta (BBA) - Molecular Cell Research, 1864(10), 1913–1920. https://doi.org/10.1016/j.bbamcr.2017.08.006

Appleby, R. D., Porteous, W. K., Hughes, G., James, A. M., Shannon, D., Wei, Y.-H., & Murphy, M. P. (1999). Quantitation and origin of the mitochondrial membrane potential in human cells lacking mitochondrial DNA. European Journal of Biochemistry, 262(1), 108–116. https://doi.org/10.1046/j.1432-1327.1999.00350.x

Atkinson, A., Smith, P., Fox, J. L., Cui, T.-Z., Khalimonchuk, O., & Winge, D. R. (2011). The LYR Protein Mzm1 Functions in the Insertion of the Rieske Fe/S Protein in Yeast Mitochondria ▿. Molecular and Cellular Biology, 31(19), 3988–3996. https://doi.org/10.1128/MCB.05673-11

Beausoleil, S. A., Villén, J., Gerber, S. A., Rush, J., & Gygi, S. P. (2006). A probability-based approach for high-throughput protein phosphorylation analysis and site localization. Nature Biotechnology, 24(10), 10. https://doi.org/10.1038/nbt1240

Becker, D., Richter, J., Tocilescu, M. A., Przedborski, S., & Voos, W. (2012). Pink1 Kinase and Its Membrane Potential (Δψ)-dependent Cleavage Product Both Localize to Outer Mitochondrial Membrane by Unique Targeting Mode. Journal of Biological Chemistry, 287(27), 22969–22987. https://doi.org/10.1074/jbc.M112.365700

Berg, J. A., Belyeu, J. R., Morgan, J. T., Ouyang, Y., Bott, A. J., Quinlan, A. R., Gertz, J., & Rutter, J. (2020). XPRESSyourself: Enhancing, standardizing, and automating ribosome profiling computational analyses yields improved insight into data. PLOS Computational Biology, 16(1), e1007625. https://doi.org/10.1371/journal.pcbi.1007625

Berg, J. A., Zhou, Y., Ouyang, Y., Waller, T. C., Cluntun, A. A., Conway, M. E., Nowinski, S. M., Van Ry, T., George, I., Cox, J. E., Wang, B., & Rutter, J. (2020). Network-aware reaction pattern recognition reveals regulatory signatures of mitochondrial dysfunction [Preprint]. Biochemistry. https://doi.org/10.1101/2020.06.25.171850

Bergwitz, C., Wee, M. J., Sinha, S., Huang, J., DeRobertis, C., Mensah, L. B., Cohen, J., Friedman, A., Kulkarni, M., Hu, Y., Vinayagam, A., Schnall-Levin, M., Berger, B., Perkins, L. A., Mohr, S. E., & Perrimon, N. (2013). Genetic Determinants of Phosphate Response in Drosophila. PLOS ONE, 8(3), e56753. https://doi.org/10.1371/journal.pone.0056753

Berry, B. J., Nieves, T. O., & Wojtovich, A. P. (n.d.). Decreased Mitochondrial Membrane Potential Activates the Mitochondrial Unfolded Protein Response. MicroPublication Biology, 2021, 10.17912/micropub.biology.000445. https://doi.org/10.17912/micropub.biology.000445

Berry, B. J., Trewin, A. J., Milliken, A. S., Baldzizhar, A., Amitrano, A. M., Lim, Y., Kim, M., & Wojtovich, A. P. (2020). Optogenetic control of mitochondrial protonmotive force to impact cellular stress resistance. EMBO Reports. https://doi.org/10.15252/embr.201949113

Berry, B. J., Vodičková, A., Müller-Eigner, A., Meng, C., Ludwig, C., Kaeberlein, M., Peleg, S., & Wojtovich, A. P. (2022). Optogenetic rejuvenation of mitochondrial membrane potential extends <EM=C. elegans</EM= lifespan. BioRxiv, 2022.05.11.491574. https://doi.org/10.1101/2022.05.11.491574

Brody, S., Oh, C., Hoja, U., & Schweizer, E. (1997). Mitochondrial acyl carrier protein is involved in lipoic acid synthesis in Saccharomyces cerevisiae. FEBS Letters, 408(2), 217–220. https://doi.org/10.1016/S0014-5793(97)00428-6

Buchet, K., & Godinot, C. (1998). Functional F1-ATPase Essential in Maintaining Growth and Membrane Potential of Human Mitochondrial DNA-depleted ρ° Cells*. Journal of Biological Chemistry, 273(36), 22983–22989. https://doi.org/10.1074/jbc.273.36.22983

Buitinck, L., Louppe, G., Blondel, M., Pedregosa, F., Mueller, A., Grisel, O., Niculae, V., Prettenhofer, P., Gramfort, A., Grobler, J., Layton, R., Vanderplas, J., Joly, A., Holt, B., & Varoquaux, G. (2013). API design for machine learning software: Experiences from the scikit-learn project. API Design for Machine Learning Software: Experiences from the Scikit-Learn Project.

Burelle, Y., Bemeur, C., Rivard, M.-E., Legault, J. T., Boucher, G., Consortium, L., Morin, C., Coderre, L., & Rosiers, C. D. (2015). Mitochondrial Vulnerability and Increased Susceptibility to Nutrient-Induced Cytotoxicity in Fibroblasts from Leigh Syndrome French Canadian Patients. PLOS ONE, 10(4), e0120767. https://doi.org/10.1371/journal.pone.0120767

Chandel, N. S. (2015). Evolution of Mitochondria as Signaling Organelles. Cell Metabolism, 22(2), 204–206. https://doi.org/10.1016/j.cmet.2015.05.013

Chen, S., Zhou, Y., Chen, Y., & Gu, J. (2018). fastp: An ultra-fast all-in-one FASTQ preprocessor. Bioinformatics, 34(17), i884–i890. https://doi.org/10.1093/bioinformatics/bty560

Chen, X. J., & Clark-Walker, G. D. (2000). The petite mutation in yeasts: 50 years on. International Review of Cytology, 194, 197–238. https://doi.org/10.1016/s0074-7696(08)62397-9

Clotet, J., Garí, E., Aldea, M., & Ariño, J. (1999). The Yeast Ser/Thr Phosphatases Sit4 and Ppz1 Play Opposite Roles in Regulation of the Cell Cycle. Molecular and Cellular Biology, 19(3), 2408–2415.

Cluntun, A. A., Badolia, R., Lettlova, S., Parnell, K. M., Shankar, T. S., Diakos, N. A., Olson, K. A., Taleb, I., Tatum, S. M., Berg, J. A., Cunningham, C. N., Van Ry, T., Bott, A. J., Krokidi, A. T., Fogarty, S., Skedros, S., Swiatek, W. I., Yu, X., Luo, B., … Drakos, S. G. (2021). The pyruvate-lactate axis modulates cardiac hypertrophy and heart failure. Cell Metabolism, 33(3), 629–648.e10. https://doi.org/10.1016/j.cmet.2020.12.003

Dasari, S., & Kölling, R. (2011). Cytosolic localization of acetohydroxyacid synthase Ilv2 and its impact on diacetyl formation during beer fermentation. Applied and Environmental Microbiology, 77(3), 727–731. https://doi.org/10.1128/AEM.01579-10

Davis, S., Weiss, M. J., Wong, J. R., Lampidis, T. J., & Chen, L. B. (1985). Mitochondrial and plasma membrane potentials cause unusual accumulation and retention of rhodamine 123 by human breast adenocarcinoma-derived MCF-7 cells. Journal of Biological Chemistry, 260(25), 13844–13850. https://doi.org/10.1016/S0021-9258(17)38802-6

Dobin, A., Davis, C. A., Schlesinger, F., Drenkow, J., Zaleski, C., Jha, S., Batut, P., Chaisson, M., & Gingeras, T. R. (2013). STAR: Ultrafast universal RNA-seq aligner. Bioinformatics, 29(1), 15–21. https://doi.org/10.1093/bioinformatics/bts635

Dupont, C.-H., Mazat, J. P., & Guerin, B. (1985). The role of adenine nucleotide translocation in the energization of the inner membrane of mitochondria isolated from ϱ+ and ϱo strains of saccharomyces cerevisiae. Biochemical and Biophysical Research Communications, 132(3), 1116–1123. https://doi.org/10.1016/0006-291X(85)91922-9

Ebrahimi, M., Habernig, L., Broeskamp, F., Aufschnaiter, A., Diessl, J., Atienza, I., Matz, S., Ruiz, F. A., & Büttner, S. (2021). Phosphate Restriction Promotes Longevity via Activation of Autophagy and the Multivesicular Body Pathway. Cells, 10(11), 3161. https://doi.org/10.3390/cells10113161

Elias, J. E., & Gygi, S. P. (2007). Target-decoy search strategy for increased confidence in large-scale protein identifications by mass spectrometry. Nature Methods, 4(3), 3. https://doi.org/10.1038/nmeth1019

Epstein, C. B., Waddle, J. A., Hale, W., Davé, V., Thornton, J., Macatee, T. L., Garner, H. R., & Butow, R. A. (2001). Genome-wide responses to mitochondrial dysfunction. Molecular Biology of the Cell, 12(2), 297–308. https://doi.org/10.1091/mbc.12.2.297

Ernst, P., Xu, N., Qu, J., Chen, H., Goldberg, M. S., Darley-Usmar, V., Zhang, J. J., O’Rourke, B., Liu, X., & Zhou, L. (2019). Precisely Control Mitochondria with Light to Manipulate Cell Fate Decision. Biophysical Journal, 117(4), 631–645. https://doi.org/10.1016/j.bpj.2019.06.038

Fernandez-Sarabia, M. J., Sutton, A., Zhong, T., & Arndt, K. T. (1992). SIT4 protein phosphatase is required for the normal accumulation of SWI4, CLN1, CLN2, and HCS26 RNAs during late G1. Genes & Development, 6(12a), 2417–2428. https://doi.org/10.1101/gad.6.12a.2417

Garipler, G., Mutlu, N., Lack, N. A., & Dunn, C. D. (2014). Deletion of conserved protein phosphatases reverses defects associated with mitochondrial DNA damage in Saccharomyces cerevisiae. Proceedings of the National Academy of Sciences, 111(4), 1473–1478. https://doi.org/10.1073/pnas.1312399111

Giaever, G., Chu, A. M., Ni, L., Connelly, C., Riles, L., Véronneau, S., Dow, S., Lucau-Danila, A., Anderson, K., André, B., Arkin, A. P., Astromoff, A., El Bakkoury, M., Bangham, R., Benito, R., Brachat, S., Campanaro, S., Curtiss, M., Davis, K., … Johnston, M. (2002). Functional profiling of the Saccharomyces cerevisiae genome. Nature, 418(6896), 387–391. https://doi.org/10.1038/nature00935

Greenleaf, A. L., Kelly, J. L., & Lehman, I. R. (1986). Yeast RPO41 gene product is required for transcription and maintenance of the mitochondrial genome. Proceedings of the National Academy of Sciences of the United States of America, 83(10), 3391–3394.

Gupta, R., Walvekar, A. S., Liang, S., Rashida, Z., Shah, P., & Laxman, S. (2019). A tRNA modification balances carbon and nitrogen metabolism by regulating phosphate homeostasis. ELife, 8, e44795. https://doi.org/10.7554/eLife.44795

Hagen, T. M., Yowe, D. L., Bartholomew, J. C., Wehr, C. M., Do, K. L., Park, J.-Y., & Ames, B. N. (1997). Mitochondrial decay in hepatocytes from old rats: Membrane potential declines, heterogeneity and oxidants increase. Proceedings of the National Academy of Sciences, 94(7), 3064–3069. https://doi.org/10.1073/pnas.94.7.3064

Heerdt, B. G., Houston, M. A., & Augenlicht, L. H. (2005). The Intrinsic Mitochondrial Membrane Potential of Colonic Carcinoma Cells Is Linked to the Probability of Tumor Progression. Cancer Research, 65(21), 9861–9867. https://doi.org/10.1158/0008-5472.CAN-05-2444

Huang, H.-M., Fowler, C., Zhang, H., & Gibson, G. E. (2004). Mitochondrial Heterogeneity Within and Between Different Cell Types. Neurochemical Research, 29(3), 651–658. https://doi.org/10.1023/B:NERE.0000014835.34495.9c

Hübscher, V., Mudholkar, K., Chiabudini, M., Fitzke, E., Wölfle, T., Pfeifer, D., Drepper, F., Warscheid, B., & Rospert, S. (2016). The Hsp70 homolog Ssb and the 14-3-3 protein Bmh1 jointly regulate transcription of glucose repressed genes in Saccharomyces cerevisiae. Nucleic Acids Research, 44(12), 5629–5645. https://doi.org/10.1093/nar/gkw168

Hughes, C. E., Coody, T. K., Jeong, M.-Y., Berg, J. A., Winge, D. R., & Hughes, A. L. (2020). Cysteine Toxicity Drives Age-Related Mitochondrial Decline by Altering Iron Homeostasis. Cell, 180(2), 296–310.e18. https://doi.org/10.1016/j.cell.2019.12.035

Hughes, C. S., Foehr, S., Garfield, D. A., Furlong, E. E., Steinmetz, L. M., & Krijgsveld, J. (2014). Ultrasensitive proteome analysis using paramagnetic bead technology. Molecular Systems Biology, 10(10), 757. https://doi.org/10.15252/msb.20145625

Hughes, C. S., Moggridge, S., Müller, T., Sorensen, P. H., Morin, G. B., & Krijgsveld, J. (2019). Single-pot, solid-phase-enhanced sample preparation for proteomics experiments. Nature Protocols, 14(1), 1. https://doi.org/10.1038/s41596-018-0082-x

Hunter, J. D. (2007). Matplotlib: A 2D Graphics Environment. Computing in Science & Engineering, 9(3), 90–95. https://doi.org/10.1109/MCSE.2007.55

Huttlin, E. L., Jedrychowski, M. P., Elias, J. E., Goswami, T., Rad, R., Beausoleil, S. A., Villén, J., Haas, W., Sowa, M. E., & Gygi, S. P. (2010). A Tissue-Specific Atlas of Mouse Protein Phosphorylation and Expression. Cell, 143(7), 1174–1189. https://doi.org/10.1016/j.cell.2010.12.001

Jablonka, W., Guzmán, S., Ramírez, J., & Montero-Lomelí, M. (2006). Deviation of carbohydrate metabolism by the SIT4 phosphatase in Saccharomyces cerevisiae. Biochimica et Biophysica Acta (BBA) - General Subjects, 1760(8), 1281–1291. https://doi.org/10.1016/j.bbagen.2006.02.014

James, A. M., Wei, Y. H., Pang, C. Y., & Murphy, M. P. (1996). Altered mitochondrial function in fibroblasts containing MELAS or MERRF mitochondrial DNA mutations. Biochemical Journal, 318(Pt 2), 401–407.

Jin, C., Barrientos, A., Epstein, C. B., Butow, R. A., & Tzagoloff, A. (2007). SIT4 regulation of Mig1p-mediated catabolite repression in Saccharomyces cerevisiae. FEBS Letters, 581(29), 5658–5663. https://doi.org/10.1016/j.febslet.2007.11.027

Jin, S. M., Lazarou, M., Wang, C., Kane, L. A., Narendra, D. P., & Youle, R. J. (2010). Mitochondrial membrane potential regulates PINK1 import and proteolytic destabilization by PARL. The Journal of Cell Biology, 191(5), 933–942. https://doi.org/10.1083/jcb.201008084

Johnson, M. A., Vidoni, S., Durigon, R., Pearce, S. F., Rorbach, J., He, J., Brea-Calvo, G., Minczuk, M., Reyes, A., Holt, I. J., & Spinazzola, A. (2014). Amino Acid Starvation Has Opposite Effects on Mitochondrial and Cytosolic Protein Synthesis. PLOS ONE, 9(4), e93597. https://doi.org/10.1371/journal.pone.0093597

Junge, W., & Nelson, N. (2015). ATP Synthase. Annual Review of Biochemistry, 84(1), 631–657. https://doi.org/10.1146/annurev-biochem-060614-034124

Kaffman, A., Herskowitz, I., Tjian, R., & O’Shea, E. K. (1994). Phosphorylation of the Transcription Factor PHO4 by a Cyclin-CDK Complex, PHO80-PHO85. Science, 263(5150), 1153–1156. https://doi.org/10.1126/science.8108735

Korshunov, S. S., Skulachev, V. P., & Starkov, A. A. (1997). High protonic potential actuates a mechanism of production of reactive oxygen species in mitochondria. FEBS Letters, 416(1), 15–18. https://doi.org/10.1016/S0014-5793(97)01159-9

Kováčová, V., Irmlerová, J., & Kováč, L. (1968). Oxidative phosphorylation in yeast. IV. Combination of a nuclear mutation affecting oxidative phosphorylation with cytoplasmic mutation to respiratory deficiency. Biochimica et Biophysica Acta (BBA) - Bioenergetics, 162(2), 157–163. https://doi.org/10.1016/0005-2728(68)90097-2

Kuro-o, M., Matsumura, Y., Aizawa, H., Kawaguchi, H., Suga, T., Utsugi, T., Ohyama, Y., Kurabayashi, M., Kaname, T., Kume, E., Iwasaki, H., Iida, A., Shiraki-Iida, T., Nishikawa, S., Nagai, R., & Nabeshima, Y. (1997). Mutation of the mouse klotho gene leads to a syndrome resembling ageing. Nature, 390(6655), 45–51. https://doi.org/10.1038/36285

Kurosu, H., Ogawa, Y., Miyoshi, M., Yamamoto, M., Nandi, A., Rosenblatt, K. P., Baum, M. G., Schiavi, S., Hu, M.-C., Moe, O. W., & Kuro-o, M. (2006). Regulation of Fibroblast Growth Factor-23 Signaling by Klotho. The Journal of Biological Chemistry, 281(10), 6120–6123. https://doi.org/10.1074/jbc.C500457200

Kurosu, H., Yamamoto, M., Clark, J. D., Pastor, J. V., Nandi, A., Gurnani, P., McGuinness, O. P., Chikuda, H., Yamaguchi, M., Kawaguchi, H., Shimomura, I., Takayama, Y., Herz, J., Kahn, C. R., Rosenblatt, K. P., & Kuro-o, M. (2005). Suppression of Aging in Mice by the Hormone Klotho. Science, 309(5742), 1829–1833. https://doi.org/10.1126/science.1112766

Lauquin, G. J. M., & Vignais, P. V. (1976). Interaction of [3H]bongkrekic acid with the mitochondrial adenine nucleotide translocator. Biochemistry, 15(11), 2316–2322. https://doi.org/10.1021/bi00656a011

Leprat, P., Ratinaud, M. H., & Julien, R. (1990). A new method for testing cell ageing using two mitochondria specific fluorescent probes. Mechanisms of Ageing and Development, 52(2), 149–167. https://doi.org/10.1016/0047-6374(90)90121-U

Li, H., Handsaker, B., Wysoker, A., Fennell, T., Ruan, J., Homer, N., Marth, G., Abecasis, G., Durbin, R., & 1000 Genome Project Data Processing Subgroup. (2009). The Sequence Alignment/Map format and SAMtools. Bioinformatics, 25(16), 2078–2079. https://doi.org/10.1093/bioinformatics/btp352

Li, J., Van Vranken, J. G., Pontano Vaites, L., Schweppe, D. K., Huttlin, E. L., Etienne, C., Nandhikonda, P., Viner, R., Robitaille, A. M., Thompson, A. H., Kuhn, K., Pike, I., Bomgarden, R. D., Rogers, J. C., Gygi, S. P., & Paulo, J. A. (2020). TMTpro reagents: A set of isobaric labeling mass tags enables simultaneous proteome-wide measurements across 16 samples. Nature Methods, 17(4), 4. https://doi.org/10.1038/s41592-020-0781-4

Liu, N.-N., Flanagan, P. R., Zeng, J., Jani, N. M., Cardenas, M. E., Moran, G. P., & Köhler, J. R. (2017). Phosphate is the third nutrient monitored by TOR in Candida albicans and provides a target for fungal-specific indirect TOR inhibition. Proceedings of the National Academy of Sciences, 114(24), 6346–6351. https://doi.org/10.1073/pnas.1617799114

Liu, S., Liu, S., He, B., Li, L., Li, L., Wang, J., Cai, T., Chen, S., & Jiang, H. (2021). OXPHOS deficiency activates global adaptation pathways to maintain mitochondrial membrane potential. EMBO Reports, 22(4), e51606. https://doi.org/10.15252/embr.202051606

Love, M. I., Huber, W., & Anders, S. (2014). Moderated estimation of fold change and dispersion for RNA-seq data with DESeq2. Genome Biology, 15(12), 550. https://doi.org/10.1186/s13059-014-0550-8

Mansell, E., Sigurdsson, V., Deltcheva, E., Brown, J., James, C., Miharada, K., Soneji, S., Larsson, J., & Enver, T. (2021). Mitochondrial Potentiation Ameliorates Age-Related Heterogeneity in Hematopoietic Stem Cell Function. Cell Stem Cell, 28(2), 241–256.e6. https://doi.org/10.1016/j.stem.2020.09.018

Martínez-Reyes, I., Diebold, L. P., Kong, H., Schieber, M., Huang, H., Hensley, C. T., Mehta, M. M., Wang, T., Santos, J. H., Woychik, R., Dufour, E., Spelbrink, J. N., Weinberg, S. E., Zhao, Y., DeBerardinis, R. J., & Chandel, N. S. (2016). TCA cycle and mitochondrial membrane potential are necessary for diverse biological functions. Molecular Cell, 61(2), 199–209. https://doi.org/10.1016/j.molcel.2015.12.002

McKinney, W. (2010). Data Structures for Statistical Computing in Python. 56–61. https://doi.org/10.25080/Majora-92bf1922-00a

Miceli, M., Jiang, J., Tiwari, A., Rodriguez-Quiñones, J., & Jazwinski, S. M. (2012). Loss of Mitochondrial Membrane Potential Triggers the Retrograde Response Extending Yeast Replicative Lifespan. Frontiers in Genetics, 2. https://www.frontiersin.org/articles/10.3389/fgene.2011.00102

Mitra, K., Wunder, C., Roysam, B., Lin, G., & Lippincott-Schwartz, J. (2009). A hyperfused mitochondrial state achieved at G1-S regulates cyclin E buildup and entry into S phase. Proceedings of the National Academy of Sciences of the United States of America, 106(29), 11960–11965. https://doi.org/10.1073/pnas.0904875106

Mouillon, J.-M., & Persson, B. L. (2006). New aspects on phosphate sensing and signalling in Saccharomyces cerevisiae. FEMS Yeast Research, 6(2), 171–176. https://doi.org/10.1111/j.1567-1364.2006.00036.x

Nowinski, S. M., Solmonson, A., Rusin, S. F., Maschek, J. A., Bensard, C. L., Fogarty, S., Jeong, M.-Y., Lettlova, S., Berg, J. A., Morgan, J. T., Ouyang, Y., Naylor, B. C., Paulo, J. A., Funai, K., Cox, J. E., Gygi, S. P., Winge, D. R., DeBerardinis, R. J., & Rutter, J. (2020). Mitochondrial fatty acid synthesis coordinates oxidative metabolism in mammalian mitochondria. ELife, 9, e58041. https://doi.org/10.7554/eLife.58041

Okuno, D., Iino, R., & Noji, H. (2011). Rotation and structure of FoF1-ATP synthase. The Journal of Biochemistry, 149(6), 655–664. https://doi.org/10.1093/jb/mvr049

Oliphant, T. (2006). *Guide to NumPy*.

Oshima, Y. (1997). The phosphatase system in Saccharomyces cerevisiae. Genes & Genetic Systems, 72(6), 323–334. https://doi.org/10.1266/ggs.72.323

Pagliarini, D. J., & Rutter, J. (2013). Hallmarks of a new era in mitochondrial biochemistry. Genes & Development, 27(24), 2615–2627. https://doi.org/10.1101/gad.229724.113

Pan, Y., Schroeder, E. A., Ocampo, A., Barrientos, A., & Shadel, G. S. (2011). Regulation of Yeast Chronological Life Span by TORC1 via Adaptive Mitochondrial ROS Signaling. Cell Metabolism, 13(6), 668–678. https://doi.org/10.1016/j.cmet.2011.03.018

Paolo, J., Magbanua, V., Ogawa, N., Harashima, S., & Oshima, Y. (1997). *The Transcriptional Activators of the PHO Regulon, Pho4p and Pho2p, Interact Directly with Each Other and with Components of the Basal Transcription Machinery in Saccharomyces cerevisiae1*.

Rad, R., Li, J., Mintseris, J., O’Connell, J., Gygi, S. P., & Schweppe, D. K. (2021). Improved Monoisotopic Mass Estimation for Deeper Proteome Coverage. Journal of Proteome Research, 20(1), 591–598. https://doi.org/10.1021/acs.jproteome.0c00563

Rohde, J. R., Campbell, S., Zurita-Martinez, S. A., Cutler, N. S., Ashe, M., & Cardenas, M. E. (2004). TOR controls transcriptional and translational programs via Sap-Sit4 protein phosphatase signaling effectors. Molecular and Cellular Biology, 24(19), 8332–8341. https://doi.org/10.1128/MCB.24.19.8332-8341.2004

Rolland, S. G., Schneid, S., Schwarz, M., Rackles, E., Fischer, C., Haeussler, S., Regmi, S. G., Yeroslaviz, A., Habermann, B., Mokranjac, D., Lambie, E., & Conradt, B. (2019). Compromised Mitochondrial Protein Import Acts as a Signal for UPRmt. Cell Reports, 28(7), 1659–1669.e5. https://doi.org/10.1016/j.celrep.2019.07.049

Sastre, J., Pallardó, F. V., Plá, R., Pellín, A., Juan, G., O’Connor, J. E., Estrela, J. M., Miquel, J., & Viña, J. (1996). Aging of the liver: Age-associated mitochondrial damage in intact hepatocytes. Hepatology, 24(5), 1199–1205. https://doi.org/10.1002/hep.510240536

Satoh, T., Enokido, Y., Aoshima, H., Uchiyama, Y., & Hatanaka, H. (1997). Changes in mitochondrial membrane potential during oxidative stress-induced apoptosis in PC12 cells. Journal of Neuroscience Research, 50(3), 413–420. https://doi.org/10.1002/(SICI)1097-4547(19971101)50:3<413::AID-JNR7>3.0.CO;2-L

Savitski, M. M., Wilhelm, M., Hahne, H., Kuster, B., & Bantscheff, M. (2015). A Scalable Approach for Protein False Discovery Rate Estimation in Large Proteomic Data Sets[S]. Molecular & Cellular Proteomics, 14(9), 2394–2404. https://doi.org/10.1074/mcp.M114.046995

Sayols, S., Scherzinger, D., & Klein, H. (2016). dupRadar: A Bioconductor package for the assessment of PCR artifacts in RNA-Seq data. BMC Bioinformatics, 17(1), 428. https://doi.org/10.1186/s12859-016-1276-2

Schindelin, J., Arganda-Carreras, I., Frise, E., Kaynig, V., Longair, M., Pietzsch, T., Preibisch, S., Rueden, C., Saalfeld, S., Schmid, B., Tinevez, J.-Y., White, D. J., Hartenstein, V., Eliceiri, K., Tomancak, P., & Cardona, A. (2012). Fiji: An open-source platform for biological-image analysis. Nature Methods, 9(7), 7. https://doi.org/10.1038/nmeth.2019

Schneider, R., Brors, B., Bürger, F., Camrath, S., & Weiss, H. (1997). Two genes of the putative mitochondrial fatty acid synthase in the genome of Saccharomyces cerevisiae. Current Genetics, 32(6), 384–388. https://doi.org/10.1007/s002940050292

Schweppe, D. K., Prasad, S., Belford, M. W., Navarrete-Perea, J., Bailey, D. J., Huguet, R., Jedrychowski, M. P., Rad, R., McAlister, G., Abbatiello, S. E., Woulters, E. R., Zabrouskov, V., Dunyach, J.-J., Paulo, J. A., & Gygi, S. P. (2019). Characterization and Optimization of Multiplexed Quantitative Analyses Using High-Field Asymmetric-Waveform Ion Mobility Mass Spectrometry. Analytical Chemistry, 91(6), 4010–4016. https://doi.org/10.1021/acs.analchem.8b05399

Sharov, V. G., Todor, A. V., Imai, M., & Sabbah, H. N. (2005). Inhibition of mitochondrial permeability transition pores by cyclosporine A improves cytochrome C oxidase function and increases rate of ATP synthesis in failing cardiomyocytes. Heart Failure Reviews, 10(4), 305–310. https://doi.org/10.1007/s10741-005-7545-1

Spinelli, J. B., & Haigis, M. C. (2018). The multifaceted contributions of mitochondria to cellular metabolism. Nature Cell Biology, 20(7), 7. https://doi.org/10.1038/s41556-018-0124-1

Sugrue, M. M., & Tatton, W. G. (2001). Mitochondrial Membrane Potential in Aging Cells. Neurosignals, 10(3–4), 176–188. https://doi.org/10.1159/000046886

Summerhayes, I. C., Lampidis, T. J., Bernal, S. D., Nadakavukaren, J. J., Nadakavukaren, K. K., Shepherd, E. L., & Chen, L. B. (1982). Unusual retention of rhodamine 123 by mitochondria in muscle and carcinoma cells. Proceedings of the National Academy of Sciences, 79(17), 5292–5296. https://doi.org/10.1073/pnas.79.17.5292

Thompson, A., Wölmer, N., Koncarevic, S., Selzer, S., Böhm, G., Legner, H., Schmid, P., Kienle, S., Penning, P., Höhle, C., Berfelde, A., Martinez-Pinna, R., Farztdinov, V., Jung, S., Kuhn, K., & Pike, I. (2019). TMTpro: Design, Synthesis, and Initial Evaluation of a Proline-Based Isobaric 16-Plex Tandem Mass Tag Reagent Set. Analytical Chemistry, 91(24), 15941–15950. https://doi.org/10.1021/acs.analchem.9b04474

Torres, J., Di Como, C. J., Herrero, E., & de la Torre-Ruiz, M. A. (2002). Regulation of the Cell Integrity Pathway by Rapamycin-sensitive TOR Function in Budding Yeast*. Journal of Biological Chemistry, 277(45), 43495–43504. https://doi.org/10.1074/jbc.M205408200

Valente, A. J., Maddalena, L. A., Robb, E. L., Moradi, F., & Stuart, J. A. (2017). A simple ImageJ macro tool for analyzing mitochondrial network morphology in mammalian cell culture. Acta Histochemica, 119(3), 315–326. https://doi.org/10.1016/j.acthis.2017.03.001

van der Walt, S., Colbert, S. C., & Varoquaux, G. (2011). The NumPy Array: A Structure for Efficient Numerical Computation. Computing in Science & Engineering, 13(2), 22–30. https://doi.org/10.1109/MCSE.2011.37

Van Vranken, J. G., Nowinski, S. M., Clowers, K. J., Jeong, M.-Y., Ouyang, Y., Berg, J. A., Gygi, J. P., Gygi, S. P., Winge, D. R., & Rutter, J. (2018). ACP Acylation Is an Acetyl-CoA-Dependent Modification Required for Electron Transport Chain Assembly. Molecular Cell, 71(4), 567–580.e4. https://doi.org/10.1016/j.molcel.2018.06.039

Vasan, K., Clutter, M., Fernandez Dunne, S., George, M. D., Luan, C.-H., Chandel, N. S., & Martínez-Reyes, I. (2022). Genes Involved in Maintaining Mitochondrial Membrane Potential Upon Electron Transport Chain Disruption. Frontiers in Cell and Developmental Biology, 10. https://www.frontiersin.org/articles/10.3389/fcell.2022.781558

Veatch, J. R., McMurray, M. A., Nelson, Z. W., & Gottschling, D. E. (2009). Mitochondrial dysfunction leads to nuclear genome instability via an iron-sulfur cluster defect. Cell, 137(7), 1247–1258. https://doi.org/10.1016/j.cell.2009.04.014

Wang, Y., & Shadel, G. S. (1999). Stability of the mitochondrial genome requires an amino-terminal domain of yeast mitochondrial RNA polymerase. Proceedings of the National Academy of Sciences, 96(14), 8046–8051. https://doi.org/10.1073/pnas.96.14.8046

Waskom M., et al. 2022. Available from: https://zenodo.org/record/7052271#.YzzMEhNKjUI

Winzeler, E. A., Shoemaker, D. D., Astromoff, A., Liang, H., Anderson, K., Andre, B., Bangham, R., Benito, R., Boeke, J. D., Bussey, H., Chu, A. M., Connelly, C., Davis, K., Dietrich, F., Dow, S. W., El Bakkoury, M., Foury, F., Friend, S. H., Gentalen, E., … Davis, R. W. (1999). Functional Characterization of the S. cerevisiae Genome by Gene Deletion and Parallel Analysis. Science, 285(5429), 901–906. https://doi.org/10.1126/science.285.5429.901

Zurita Rendón, O., Fredrickson, E. K., Howard, C. J., Van Vranken, J., Fogarty, S., Tolley, N. D., Kalia, R., Osuna, B. A., Shen, P. S., Hill, C. P., Frost, A., & Rutter, J. (2018). Vms1p is a release factor for the ribosome-associated quality control complex. Nature Communications, 9(1), 1. https://doi.org/10.1038/s41467-018-04564-3

